# WRNIP1-ATM signaling promotes G-quadruplex resolution by preserving FANCJ stability

**DOI:** 10.64898/2026.07.02.736033

**Authors:** Pasquale Valenzisi, Rosa Parrillo, Pietro Pichierri, Annapaola Franchitto

## Abstract

G-quadruplexes (G4s) are non-canonical DNA structures that regulate transcription, replication, and DNA repair but, when unresolved, hinder replication fork progression and compromise genome stability.

Here, we identify a WRNIP1-dependent mechanism that promotes the resolution of R-loop-associated G4 structures. Loss of WRNIP1 or disruption of its ubiquitin-binding zinc finger (UBZ) domain causes persistent G4/R-loop accumulation, leading to transcription-replication conflicts, DNA damage, and genome instability.

We further show that WRNIP1 interacts with the G4 helicase FANCJ and that mutation of the UBZ domain disrupts this interaction, impairing FANCJ stability and recruitment to G4 sites. Mechanistically, WRNIP1 acts upstream of FANCJ by promoting ATM-dependent signaling required for FANCJ stabilization and chromatin association. ATM-mediated phosphorylation of FANCJ at Ser990 protects the helicase from ubiquitin-dependent proteasomal degradation.

Together, these findings define a WRNIP1-ATM-FANCJ regulatory axis that promotes G4 resolution during DNA replication and preserves genome stability.

## INTRODUCTION

DNA replication is a tightly regulated process in living cells, essential for ensuring the accurate duplication of genetic material. However, it constantly faces several challenges that can hinder replication and potentially lead to errors or mutations in the newly synthesized DNA. One such challenge is posed by non-canonical DNA secondary structures ^1^. The human genome can adopt several non-canonical conformations, including DNA hairpins, Holliday junctions, cruciforms, triplexes, R-loops, and G-quadruplexes (G4s) ^2,3,4^. Unresolved DNA secondary structures, in general, and G4s in particular, are major sources of replication stress and genomic instability ^5^.

G4s are stable DNA structures formed by stacked guanine tetrads held together by Hoogsteen hydrogen bonds. They spontaneously assemble on G-rich single-stranded DNA (ssDNA) exposed by the advancing replication fork, potentially causing the uncoupling of replication components and resulting in replication fork stalling ^4^. G4s are not randomly distributed throughout the genome and regulate essential cellular processes, including DNA replication, transcription, recombination, and gene expression ^6,7^.

In recent years, the study of G4s has gained increasing attention due to their involvement in malignant transformation and cancer development, as well as their potential as promising targets for anticancer therapy ^8^. An intriguing interplay between G4s and R-loops, three-stranded nucleic acid structures consisting of DNA-RNA hybrids and ssDNA displaced from the non-template strand, has been observed in cancer cells ^9^. It has been suggested that G-rich sequences within the non-template strand of R-loops can form a G4 motif that contribute to stabilizing the R-loop itself ^10^. In eukaryotic cells, the persistence of G4/R-loops can lead to harmful interference between the replication and transcription machineries, known as transcription-replication conflicts (TRCs), thereby inducing replication stress, a hallmark of pre-cancerous and cancerous cells ^11^.

G4s pose significant challenges to DNA replication, requiring specialized DNA helicases to promote their unwinding and ensure smooth DNA synthesis ^12,13^. While several helicases have been shown to efficiently resolve G4 structures *in vitro*, their precise functions *in vivo* remain only partially defined^14^.

Among these factors, FANCJ, mutated in hereditary breast and ovarian cancer, as well as in the chromosomal instability disorder Fanconi anaemia, is considered one of the most potent G4 resolvases during DNA synthesis ^15,16^. Recent evidence further suggests that FANCJ, together with MutSβ and MutLβ, contributes to the removal of G4/R-loops at TRC sites, facilitating replication restart ^17^. Nevertheless, the molecular mechanisms underlying G4 removal at R-loop-mediated TRCs remain poorly understood. Elucidating these processes could provide important insights into how cells preserve replication fork progression and genome stability.

The fork-protective factor WRN helicase-interacting protein 1 (WRNIP1) belongs to the highly conserved AAA+ family of ATPases and also possesses a ubiquitin-binding zinc finger (UBZ) domain. It plays a crucial role in various DNA-related processes, including replication, repair, and recombination, significantly contributing to the maintenance of genome stability ^18^. WRNIP1 is recruited to stalled replication forks, where it interacts with RAD51 and BRCA2 ^19^. This interaction is essential for protecting stalled forks from uncontrolled MRE11-mediated degradation, preventing replication stress. The ATPase activity of WRNIP1 is specifically required for the restart of stalled forks ^19^.

Under specific pathological conditions, WRNIP1 remains bound to chromatin to enhance RAD51 interaction with ssDNA near R-loops ^20^. A direct involvement of the UBZ domain of WRNIP1 in counteracting R-loop persistence following mild replication perturbation has been demonstrated ^21^. The close proximity of WRNIP1 to transcription/replication complexes and R-loops following replication perturbation suggests its active role in resolving TRCs and thereby averting genomic instability ^21^.

The yeast homolog of WRNIP1, Mgs1 (maintenance of genome stability), has been identified as a G4-binding protein ^22,23^. Mgs1 is thought to support replication at G4-forming regions by protecting stalled replication forks and avoiding genomic instability ^23^. WRNIP1 has also been reported to exhibit robust *in vitro* G4 DNA binding affinity ^24^. However, the precise mechanism by which WRNIP1 deals with G4s in cells remains unclear.

This work identifies the WRNIP1-ATM-FANCJ axis as a previously unappreciated mechanism that safeguards genome stability by facilitating the resolution of G-quadruplex structures during DNA replication. In addition, our findings elucidate how cells respond to G4-associated replication stress and highlight the central role of WRNIP1-mediated ATM signalling in regulating FANCJ helicase activity to maintain genome stability.

## RESULTS

### WRNIP1 loss or UBZ mutation accumulates G4 at R-loops

The G-rich sequences on the non-template strand of R-loops can fold into G-quadruplexes (G4) structures, which stabilize the RNA-DNA hybrid and reinforce R-loop persistence, potentially compromising genome stability ^9,25,4^. We previously showed that loss of WRNIP1 or mutation in its ubiquitin-binding zinc finger (UBZ) domain promotes R-loop accumulation under replication stress ^21^. To determine whether this accumulation depends on G4 stabilization, we measured G4 levels in cells exposed to low-dose aphidicolin (Aph) using the G4-specific BG4 antibody ^26^.

WRNIP1-deficient (shWRNIP1) and UBZ mutant (shWRNIP1^D37A^) cells presented a marked increase in nuclear BG4 signal compared to wild-type (shWRNIP1^WT^) and ATPase mutant (shWRNIP1^T294A^) cells (Figure 1A). This effect was further amplified by Aph treatment specifically in WRNIP1-deficient and UBZ mutant cells (Figure 1A). Importantly, transcription inhibition or RNase H1 overexpression, known to suppress R-loops ^27^, significantly reduced BG4 staining (Figure 1A and B), indicating that the presence of G4 structures is closely linked to R-loop formation. Consistent results were obtained using low doses of HU and CPT, indicating that the effect is not drug-specific (Supplementary Figure 1A).

**Figure 1.**
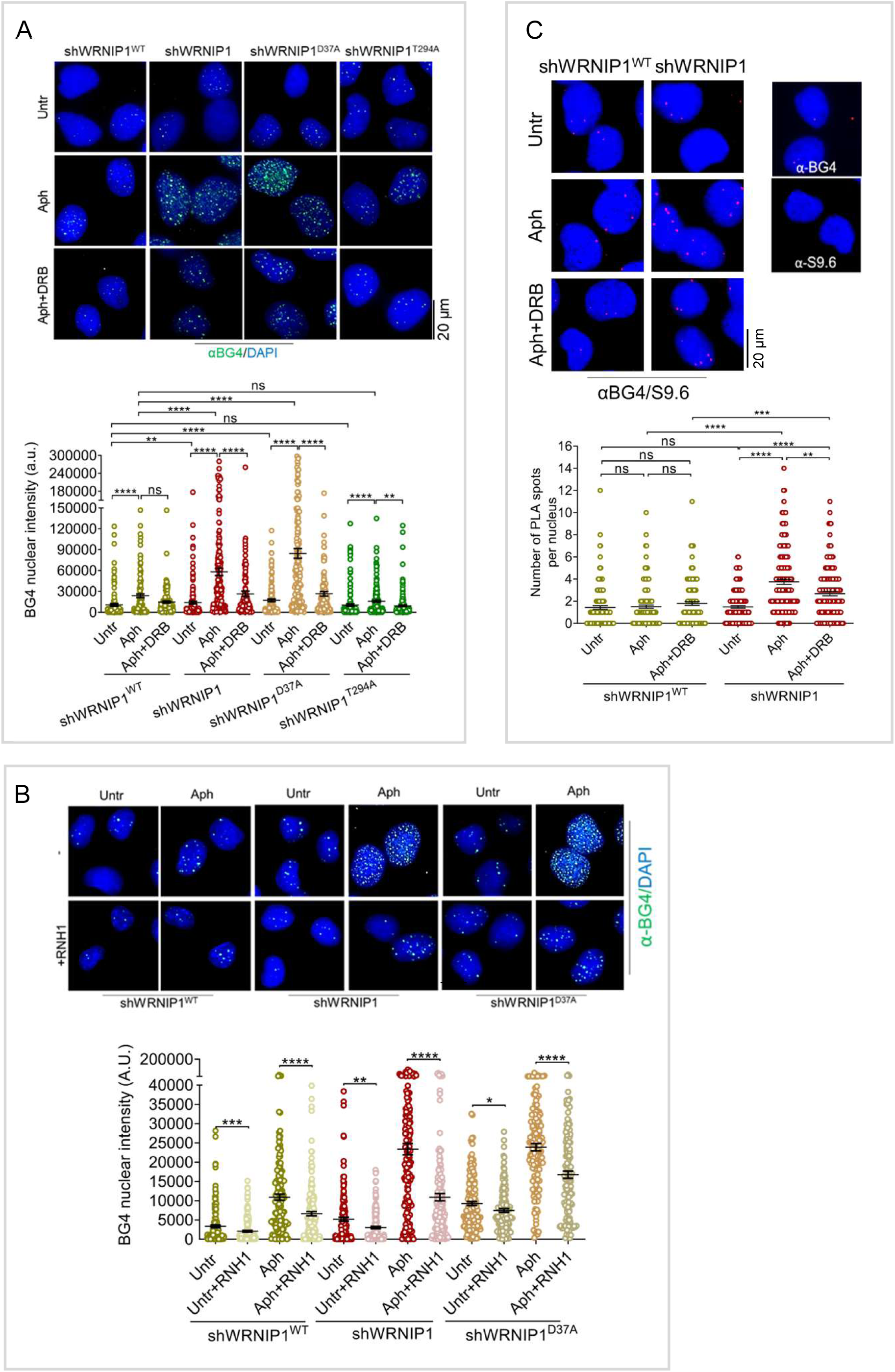
G4/R-loops accumulate in WRNIP1-deficient or UBZ mutant cells. (A) **Immunofluorescence analysis of G4 accumulation.** shWRNIP1^WT^, shWRNIP1, shWRNIP1^D37A^ and shWRNIP1^T294A^ cells were treated or not with Aph for 24 h, with DRB added during the last 3 h, fixed and stained with BG4 (G4s) antibody. DNA was counterstained with DAPI. Representative images are shown. Dot plot shows nuclear BG4 fluorescence intensity. Horizontal lines indicate median values (ns: *P* > 0.05; \*\**P* < 0.01; \*\*\*\**P* < 0.0001; Mann-Whitney test). (B) **G4 levels assessed by immunofluorescence.** shWRNIP1^WT^, shWRNIP1 and shWRNIP1^D37A^ cells were transfected with GFP-tagged RNaseH1 or empty vector and, 48 h later, treated or not with Aph for 24 h, fixed and stained with BG4 (G4s) antibody. DNA was counterstained with DAPI. Representative images are shown. Dot plot shows nuclear BG4 fluorescence intensity. Horizontal lines indicate the median values (\**P* < 0.05, \*\**P* < 0.01; \*\*\**P* < 0.001; \*\*\*\**P* < 0.0001; Mann-Whitney test). (C) **G4-R-loop interactions detected by PLA.** shWRNIP1^WT^ and shWRNIP1 cells treated or not with Aph for 24 h, alone or in combination with DRB during the last 3 h, fixed and subjected to PLA using BG4 (G4s) and the S9.6 (R-loops) antibodies. DNA was counterstained with DAPI. Representative images are shown. Dot plot shows the number of PLA spots per nucleus. Horizontal lines indicate the median values (ns: *P* > 0.05; \*\**P* < 0.01; \*\*\**P* < 0.001; \*\*\*\**P* < 0.0001; Kruskal-Wallis test).

Proximity ligation assays (PLA) confirmed the close spatial proximity between G4s and R-loops in WRNIP1-deficient cells after Aph treatment, an effect attenuated by transcription inhibition (Figure 1C). The G4-stabilizing ligand pyridostatin (PDS) ^28^ similarly increased BG4 signal and selectively impaired viability of WRNIP1-deficient and UBZ mutant cells (Supplementary Figure 1B and C). Conversely, high doses Aph, HU and CPT, which arrest replication, suppressed G4 accumulation (Supplementary Figure 1A and D), suggesting that G4 formation is largely linked to ongoing DNA synthesis. Consistently, EdU (5-ethynyl-2’-deoxyuridine)-positive WRNIP1-deficient and UBZ mutant cells, representing cells actively undergoing DNA synthesis, exhibited higher G4 accumulation following Aph treatment compared to EdU-negative cells, supporting that G4 formation predominantly occurs during S phase (Supplementary Figure 1E).

Next, to demonstrate that G4s form on R-loops, cells were pre-labelled with the thymidine analogue 5-iodo-2’-deoxyuridine (IdU) to mark the parental ssDNA, including the displaced ssDNA strand of R-loops, and then treated with Aph (Figure 2A). In WRNIP1-deficient cells, Aph treatment led to an increase in ssDNA signal, which was reduced upon transcription inhibition (Figure 2A).

**Figure 2.**
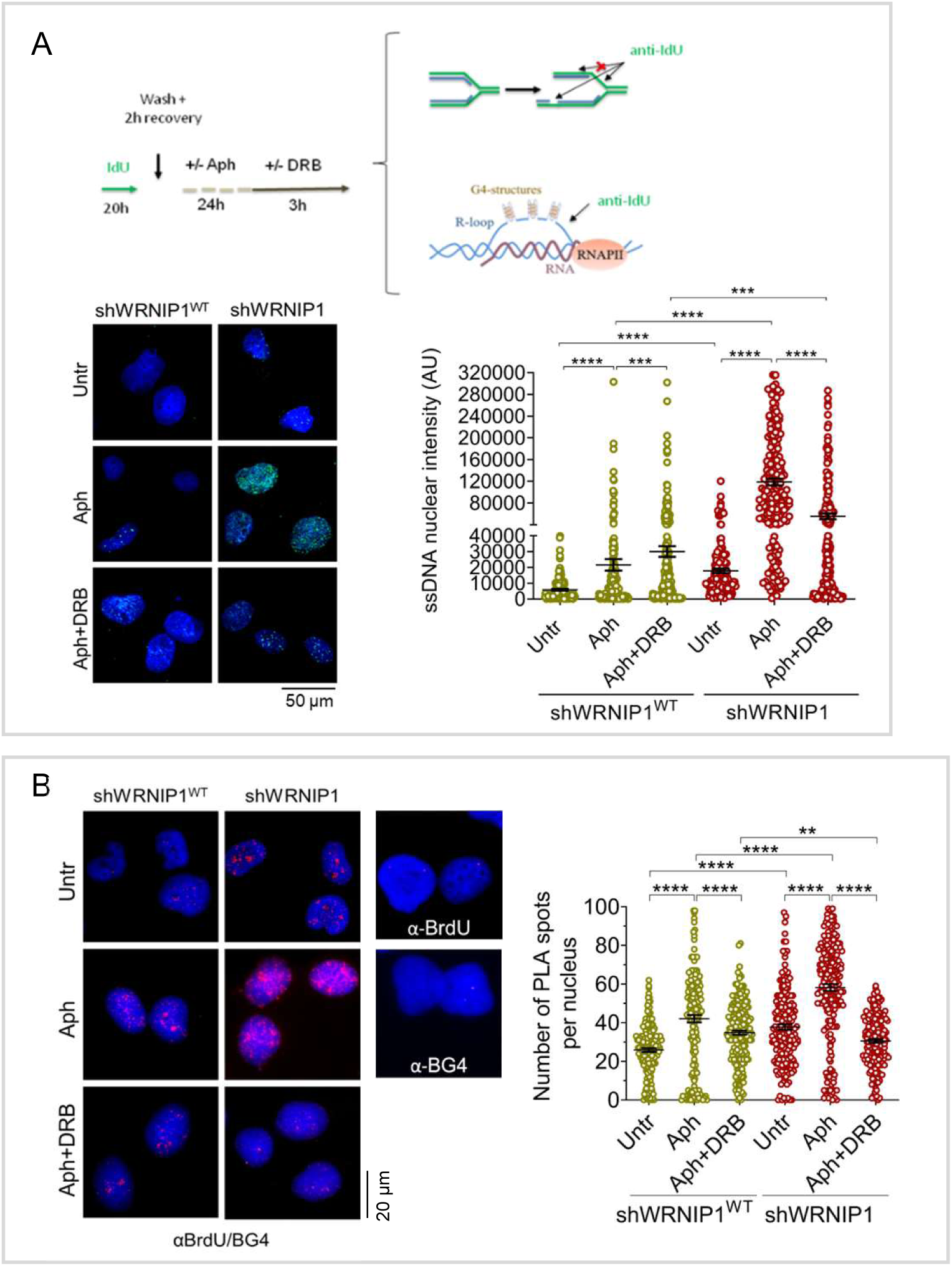
G4s co-localize with R-loop-displaced ssDNA. (A) **Parental ssDNA strand accumulation.** shWRNIP1^WT^ and shWRNIP1 cells were treated as indicated in the experimental scheme, fixed and stained with an anti-IdU antibody to detect parental ssDNA strand. DNA was counterstained with DAPI. Representative images are shown. Dot plot shows nuclear IdU fluorescence intensity. Horizontal lines indicate median values (\*\*\**P* < 0.001; \*\*\*\**P* < 0.0001; Mann-Whitney test). (B) **G4-parental ssDNA strand interaction detected by PLA.** shWRNIP1^WT^ and shWRNIP1 cells treated as in (A) and subjected to PLA using BG4 (G4s) antibody and anti-IdU (ssDNA) antibodies. DNA was counterstained with DAPI. Representative images are shown. Dot plot shows the number of PLA spots per nucleus. Horizontal lines indicate median values (\*\**P* < 0.01; \*\*\*\**P* < 0.0001; Kruskal-Wallis test).

Enhanced proximal localization between ssDNA (anti-IdU) and G4 structures (anti-BG4) was observed following Aph treatment in both wild-type and WRNIP1-deficient cells, with a more pronounced effect in WRNIP1-deficient cells (Figure 2B). Notably, this proximal localization was strongly reduced by DRB. Thus, consistent with the increased association between G4s and R-loops (Figure 1C), this suggests that most ssDNA is proximal to R-loops.

Together, these findings demonstrate that loss of WRNIP1 or mutation in its UBZ domain promotes R-loop-associated G4 accumulation.

### Persistent G4/R-loops drive TRCs and impair DNA repair

G4-mediated stabilization of co-transcriptional R-loops can trigger transcription-replication conflicts (TRCs) ^29^. Consistent with this, loss of WRNIP1 or mutation in its UBZ domain results in R-loop-dependent TRCs ^21^, as well as increased G4 accumulation and enhanced spatial proximity between R-loops and G4s (Figure 1B and C). Treatment with PDS exacerbated TRCs in WRNIP1-deficient and UBZ mutant cells compared to wild-type cells, whereas pre-treatment with high doses replication (Aph) and transcription (DRB) inhibitors, which prevent TRC formation ^30^, restored TRC levels and blocked G4 formation (Figure 3A; Supplementary Figure 2A), demonstrating that G4/R-loop accumulation drives TRC formation.

**Figure 3.**
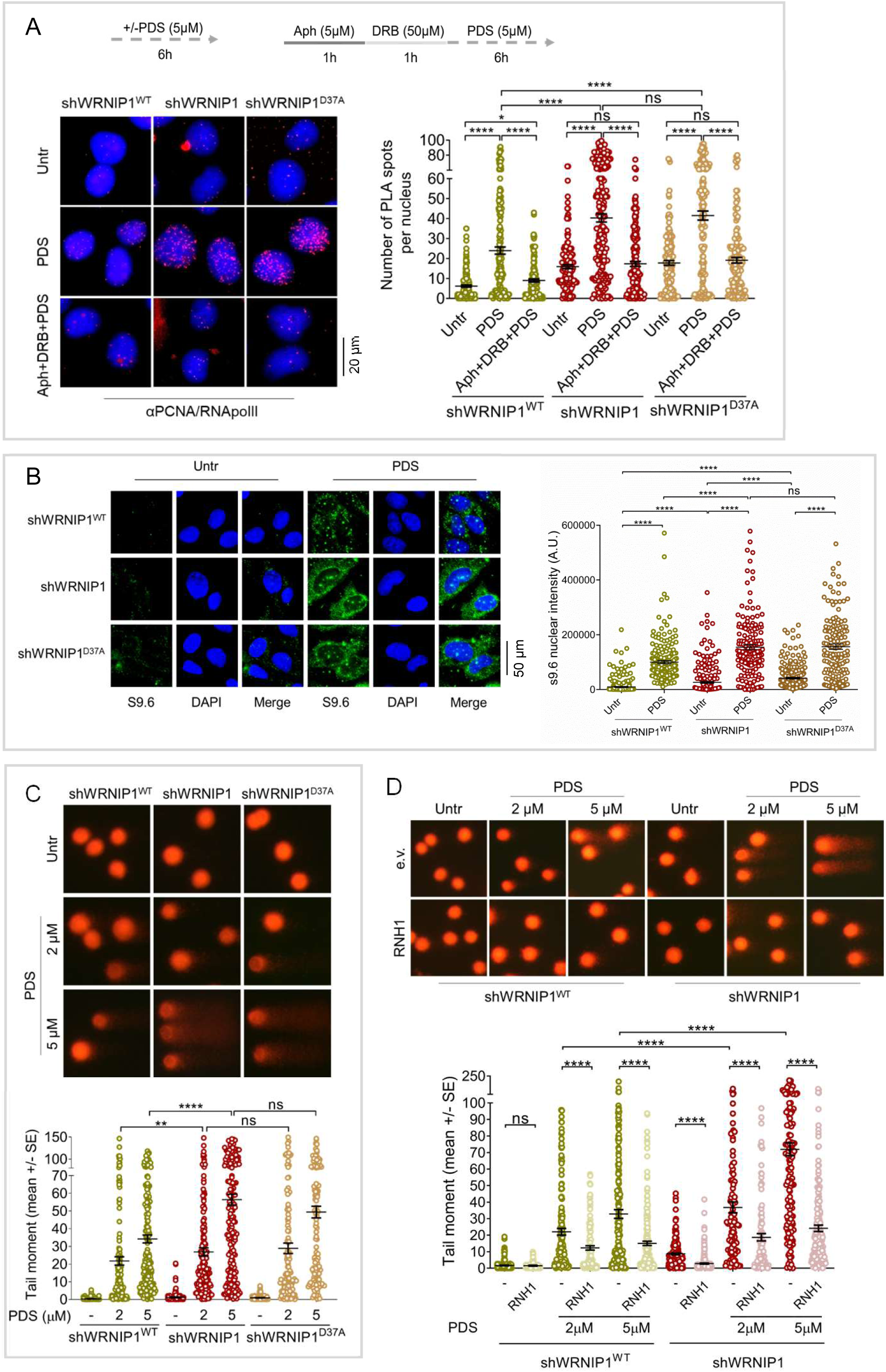
G4/R-loop-dependent DNA damage accumulation in WRNIP1-deficient or UBZ mutant cells. (A) **Detection of TRCs by PLA.** shWRNIP1^WT^, shWRNIP1 and shWRNIP1^D37A^ cells were treated as indicated in the experimental scheme, fixed and subjected to PLA using PCNA and RNA polymerase II (RNA polII) antibodies. DNA was counterstained with DAPI. Representative images are shown. Dot plot shows the number of PLA spots per nucleus. Horizontal lines indicate median values (ns: *P* > 0.05; \**P* < 0.05; \*\*\*\**P* < 0.0001; Kruskal-Wallis test). (B) **R-loop accumulation after prolonged PDS exposure.** shWRNIP1^WT^, shWRNIP1 and shWRNIP1^D37A^ cells were treated or not with 5 µM PDS for 24 h, fixed and stained with S9.6 antibody to detect R-loops. DNA was counterstained with DAPI. Representative images are shown. Dot plot shows nuclear S9.6 fluorescence intensity. Horizontal lines indicate median values (ns: *P* > 0.05; \*\*\*\**P* < 0.0001; Mann-Whitney test). (C) **PDS-induced DNA damage assessed by alkaline Comet assay.** shWRNIP1^WT^and shWRNIP1 cells were treated or not with 2 or 5 µM PDS for 24 h. Representative images are shown. Dot plot shows tail moment values. Horizontal lines indicate median values (ns: *P* > 0.05; \*\**P* < 0.01; \*\*\*\**P* < 0.0001; Mann-Whitney test). (D) **Rescue of DNA damage by RNaseH1 overexpression assessed through alkaline Comet assay.** shWRNIP1^WT^ and shWRNIP1 cells were transfected with GFP-tagged RNaseH1 (RNH1) or empty vector and treated or not with 2 or 5 µM PDS for 24 h. Representative images are shown. Dot plot shows mean tail moment. Horizontal lines indicate median values (ns: *P* > 0.05; \*\*\*\**P* < 0.0001; Mann-Whitney test).

G4 stabilization also increased nuclear R-loops following short- and long-term PDS exposure, particularly in WRNIP1-deficient and UBZ mutant cells (Figure 3B; Supplementary Figure 2B), consistent with prior report ^9^. Functionally, alkaline Comet assay revealed elevated spontaneous and PDS-induced DNA damage in WRNIP1-deficient and UBZ mutant cells, accompanied by increased double-strand breaks (DSBs), as previously demonstrated ^30^ (Figure 3C; Supplementary Figure 3A). RNase H1 overexpression and replication/transcription inhibition markedly reduced DNA damage and DSB formation, confirming that G4/R-loop accumulation is a key driver of genome instability in the absence of functional WRNIP1 (Figure 3D; Supplementary Figure 3B and C).

We next assessed DNA repair capacity. After Aph release, alkaline Comet assay showed that wild-type cells repaired DNA damage within 3 hours, whereas WRNIP1-deficient cells retained high damage levels (Figure 4A). In contrast, cells lacking ATPase activity, but not UBZ mutant, recovered after 6 hours (Figure 4B). Neutral Comet assay confirmed the presence of DSBs in WRNIP1-deficient and UBZ mutant cells (Supplementary Figure 4). These findings indicate that G4/R-loop accumulation in WRNIP1-deficient and UBZ mutant cells drives TRCs and delays DNA repair.

**Figure 4.**
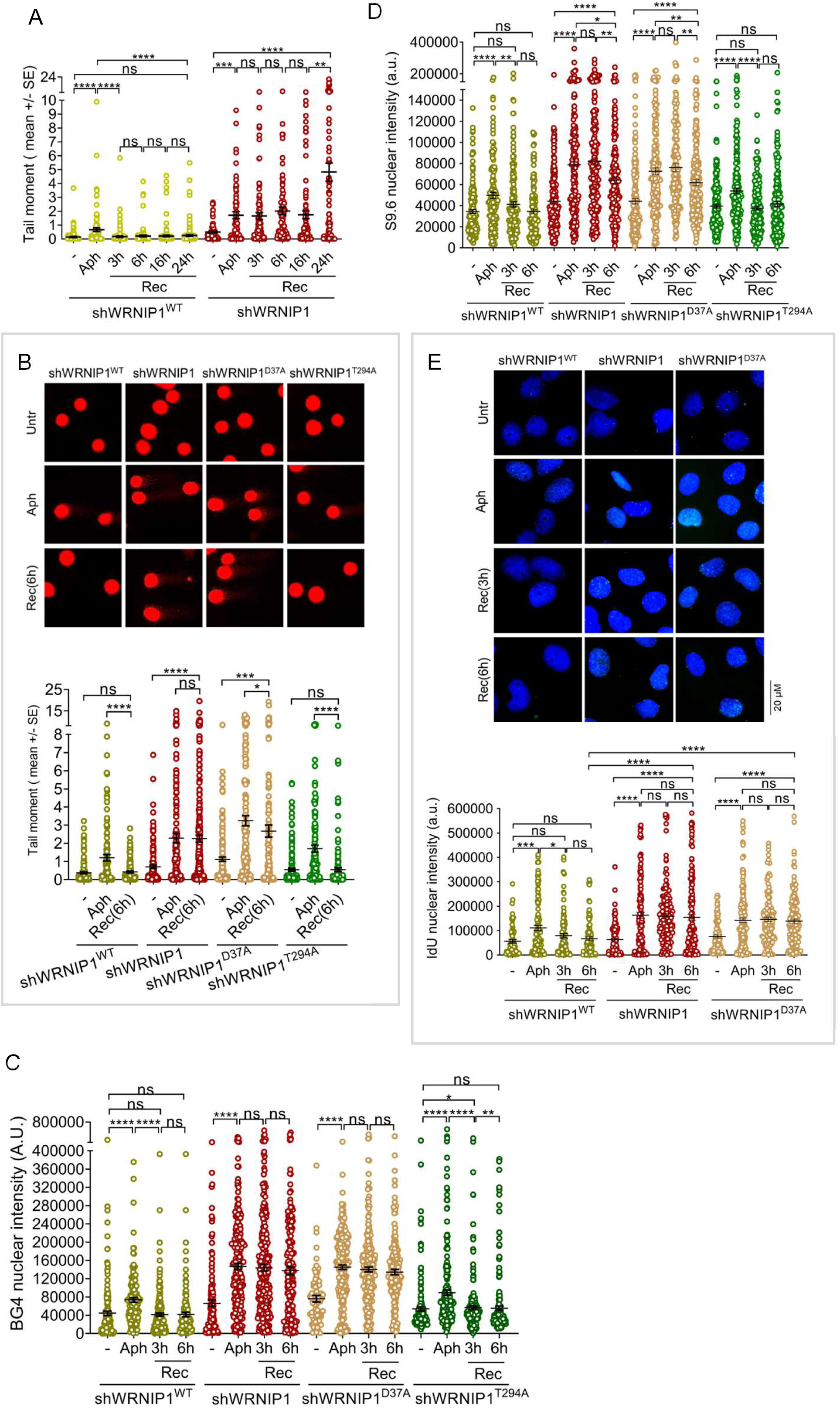
Persistent G4s increase DNA damage in WRNIP1-deficient and UBZ mutant cells. (A) **Recovery from Aph-induced damage assessed by alkaline Comet assay.** shWRNIP1^WT^ and shWRNIP1 cells were treated with or without Aph for 24 h and allowed to recover in drug-free medium for the indicated times. Dot plot shows tail moment values. Horizontal lines indicate median values (ns: *P* > 0.05; \*\**P* < 0.01; \*\*\**P* < 0.001; \*\*\*\**P* < 0.0001; Mann-Whitney test). (B) **Impact of WRNIP1 loss or mutation on recovery from Aph-induced DNA damage**. shWRNIP1^WT^, shWRNIP1, shWRNIP1^D37A^ and shWRNIP1^T294A^ cells were treated as in (A) and analysed by alkaline Comet assay. Representative images are shown. Dot plot shows tail moment values. Horizontal lines indicate median values (ns: *P* > 0.05; \**P* < 0.05; \*\*\**P* < 0.001; \*\*\*\**P* < 0.0001; Mann-Whitney test). (C) **Immunofluorescence analysis of G4 levels.** Cells were treated as in (A), fixed and stained with BG4 (G4s) antibody. DNA was counterstained with DAPI. Dot plot shows nuclear BG4 fluorescence intensity. Horizontal lines indicate median values (ns: *P* > 0.05; \**P* < 0.05; \*\**P* < 0.01; \*\*\*\**P* < 0.0001; Mann-Whitney test). (D) **R-loop accumulation following recovery from Aph exposure.** Cells were treated as in (A), allowed to recover in drug-free medium for the indicated times, fixed and stained with S9.6 antibody to detect R-loops. DNA was counterstained with DAPI. Dot plot shows nuclear S9.6 fluorescence intensity. Horizontal lines indicate median values ((ns: *P* > 0.05; \**P* < 0.05; \*\**P* < 0.01; \*\*\*\**P* < 0.0001; Mann-Whitney test). (E) **Accumulation of parental ssDNA strand following recovery from Aph.** shWRNIP1^WT^, shWRNIP1 and shWRNIP1^D37A^ cells were treated as in (A), allowed to recover in drug-free medium for the indicated times, fixed and stained with anti-IdU antibody to detect parental ssDNA strand. DNA was counterstained with DAPI. Representative images are shown. Dot plot shows nuclear IdU fluorescence intensity. Horizontal lines indicate median values (ns: *P* > 0.05; \**P* < 0.05; \*\*\**P* < 0.001; \*\*\*\**P* < 0.0001; Mann-Whitney test).

We hypothesized that the observed repair defect observed in WRNIP1-deficient and UBZ mutant cells results from persistent G4 structures. Supporting this, BG4 staining revealed increased nuclear G4 intensity after Aph treatment, which remained high at 3 and 6 hours of recovery in WRNIP1-deficient and UBZ mutant cells, but not in ATPase mutant or wild-type cells (Figure 4C). R-loop levels, which can associate with G4s declined in wild-type and ATPase mutant cells by 3 hours and returned to baseline by 6 hours, with only a modest reduction at 6 hours (Figure 4D), indicating impaired resolution of G4-associated R-loops.

To further examine the persistence of displaced ssDNA associated with R-loops, cells were pre-labelled with IdU and allowed to recover in drug-free medium. Immunofluorescence detection showed a time-dependent decrease in IdU signal in wild-type cells, whereas IdU intensity remained largely unchanged in WRNIP1-deficient and UBZ mutant cells, suggesting the presence of unresolved ssDNA at or near G4 structures (Figure 4E).

Overall, loss of WRNIP1 or mutation in its UBZ domain leads to persistence of G4 and R-loop structures, delaying DNA repair. In contrast, ATPase-mutant cells efficiently resolve these structures, highlighting the critical role of the UBZ domain in maintaining genome integrity.

### WRNIP1 deficiency and UBZ domain mutation destabilize FANCJ

Mgs1, the yeast homologue of WRNIP1, binds G4 DNA to facilitate replication at G4-forming regions and preserve genome integrity ^23^. Similarly, WRNIP1 exhibits strong *in vitro* affinity for G4 DNA ^24^. To determine whether WRNIP1 associates with G4s, we performed PLA analysis.

Consistent with yeast data ^23^, WRNIP1 was found in close proximity to G4s at comparable levels in wild-type and ATPase mutant cells, under both normal and Aph-treated conditions, but increased in UBZ mutant cells, likely due to higher G4 abundance (Figure 5A).

**Figure 5.**
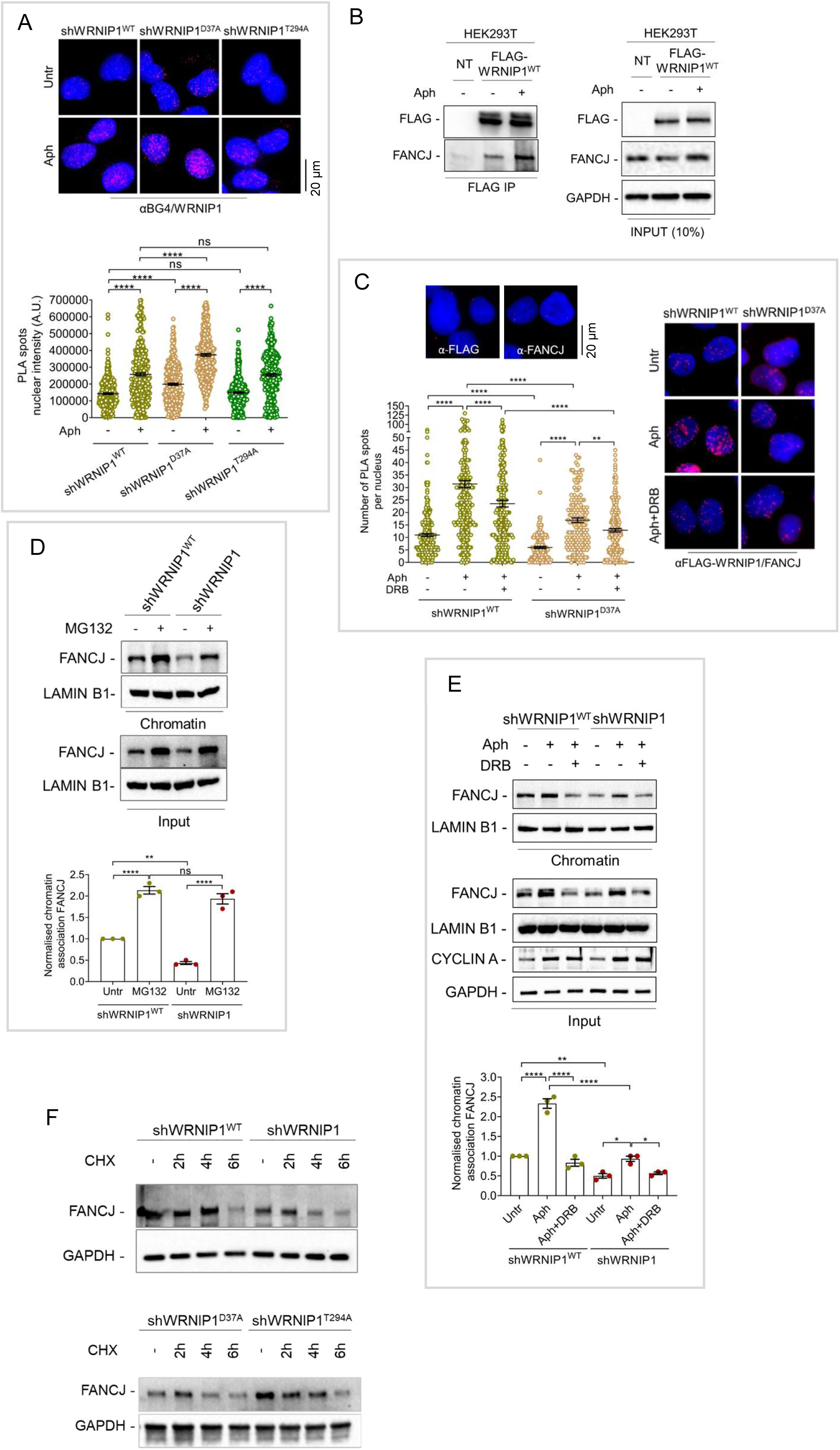
WRNIP1 and FANCJ interact upon G4/R-loop accumulation. (A) **G4-WRNIP1 interactions detected by PLA.** shWRNIP1^WT^, shWRNIP1^D37A^ and shWRNIP1^T294A^ cells were treated or not with Aph for 24 h, fixed and subjected to PLA using BG4 (G4s) and anti-Flag antibodies. DNA was counterstained with DAPI. Representative images are shown. Dot plot shows the nuclear BG4 fluorescence intensity. Horizontal lines indicate median values (ns: *P* > 0.05; \*\*\*\**P* < 0.0001; ANOVA test). (B) **Co-immunoprecipitation of WRNIP1 and FANCJ.** HEK293T cells were transfected with FLAG-WRNIP1^WT^ and treated or not with 0.4 µM Aph for 24 h. IP was performed with anti-FLAG antibody. Blots were probed with antibodies against WRNIP1, FLAG and GAPDH. (C) **WRNIP1-FANCJ interactions detected by PLA.** shWRNIP1^WT^ and shWRNIP1^D37A^ cells were treated or not with Aph for 24 h and DRB during the last 3 h, fixed and subjected to PLA using anti-FLAG (WRNIP1) and anti-FANCJ antibodies. DNA was counterstained with DAPI. Representative images are shown. Dot plot shows the number of PLA spots per nucleus. Horizontal lines indicate median values (\*\**P* < 0.01; \*\*\*\**P* < 0.0001; Kruskal-Wallis test). (D) **FANCJ association with chromatin.** shWRNIP1^WT^ and shWRNIP1 cells were treated or not with MG132 for 4 h. Chromatin fractions were immunoblotted with antibodies against FANCJ and LAMIN B1. Data are shown as super plot of FANCJ/LAMIN B1 ratios normalized to control (ns: *P* > 0.05; \*\**P* < 0.01; \*\*\*\**P* < 0.0001; one-way ANOVA test). (E) **FANCJ association with chromatin following transcription inhibition.** shWRNIP1^WT^ and shWRNIP1 cells were treated or not with Aph for 24 h and DRB during the last 3 h. Blots were probed with antibodies against FANCJ, LAMIN B1, CYCLIN A and GAPDH. CYCLIN A marks S-phase cells. Data are shown as super plot of FANCJ/GAPDH ratios normalized to control (\**P* < 0.05; \*\**P* < 0.01; \*\*\*\**P* < 0.0001; one-way ANOVA test). (F) **Analysis of FANCJ stability**. shWRNIP1^WT^, shWRNIP1, shWRNIP1^D37A^ and shWRNIP1^T294A^ cells were treated or not with 40 µg/mL cycloheximide as indicated to assess FANCJ stability. Blots were probed with antibodies against FANCJ and GAPDH.

In human cells, FANCJ is one of the major DNA helicases responsible for unwinding G4 structures ^13^. To explore a potential functional link between WRNIP1 and FANCJ, we performed co-immunoprecipitation. WRNIP1 and FANCJ interacted under untreated conditions, with their association increasing upon Aph treatment (Figure 5B). Immunoprecipitation was also performed using FANCJ as the bait protein, confirming the interaction under the same conditions (Supplementary Figure 5A). PLA experiments corroborated these results, showing that Aph treatment enhanced the spatial proximity between WRNIP1 and FANCJ, which was reduced by transcription inhibition, and that UBZ mutation impaired this association (Figure 5C).

Loss of WRNIP1 or UBZ domain mutation reduced both total and chromatin-bound FANCJ levels (Figure 5D and E; Supplementary Figure 5B). Notably, although proteasome inhibition restored FANCJ levels in both cell lines, the baseline levels in shWRNIP1 cells were lower (Figure 5D). These data indicate that WRNIP1 protects FANCJ from ubiquitin-mediated degradation. Moreover, DRB treatment reduced both total and chromatin-bound FANCJ levels, likely due to decreased G4 accumulation (Figure 5E). Mutation of the ATPase domain had no effect, reinforcing specific role of the UBZ domain in FANCJ stabilization (Supplementary Figure 5B). Similar results were observed after treatment with the G4 stabilizer PDS (Supplementary Figure 5C), and defective FANCJ stabilization in WRNIP1-deficient cells was consistent across different cell types (Supplementary Figure 5D).

Finally, to confirm proteasome-mediated degradation, wild-type, WRNIP1-deficient, and UBZ mutant cells were treated with cycloheximide and protein stability was monitored by Western blotting. FANCJ degraded more rapidly in WRNIP1-deficient cells than in wild-type cells, whereas co-treatment with MG132 restored stability (Figure 5F and Supplementary Figure 5E).

These findings demonstrate that WRNIP1 interacts with FANCJ and protects it from degradation, with the UBZ domain playing a critical role in maintaining FANCJ stability under replication stress.

### WRNIP1 and FANCJ define a common G4 resolution pathway

Previous study showed that G4-stabilizing ligands increase cellular dependence on the DNA helicase FANCJ ^31^. In line with this, FANCJ depletion leads to G4 accumulation, DNA damage, and persistent replication fork stalling at G4 structures ^32^, emphasizing its essential role in G4 unwinding. To understand how WRNIP1 contributes to this process, we examined the combined effects of WRNIP1 loss or UBZ mutation with FANCJ depletion.

We first measured G4 levels following Aph treatment in FANCJ-depleted cells. Immunofluorescence analysis showed that loss of either WRNIP1 or FANCJ significantly increased BG4 nuclear intensity, which was further enhanced upon Aph treatment (Figure 6A). Notably, simultaneous loss of WRNIP1 and FANCJ did not further increase G4 levels compared to single deficiencies (Figure 6A; Supplementary Figure 6A). Consistently, FANCJ loss induced similar levels of Aph-associated DNA damage in both wild-type and WRNIP1-depleted cells (Supplementary Figure 6B). These results suggest that WRNIP1 and FANCJ act in the same pathway to limit G4 accumulation.

**Figure 6.**
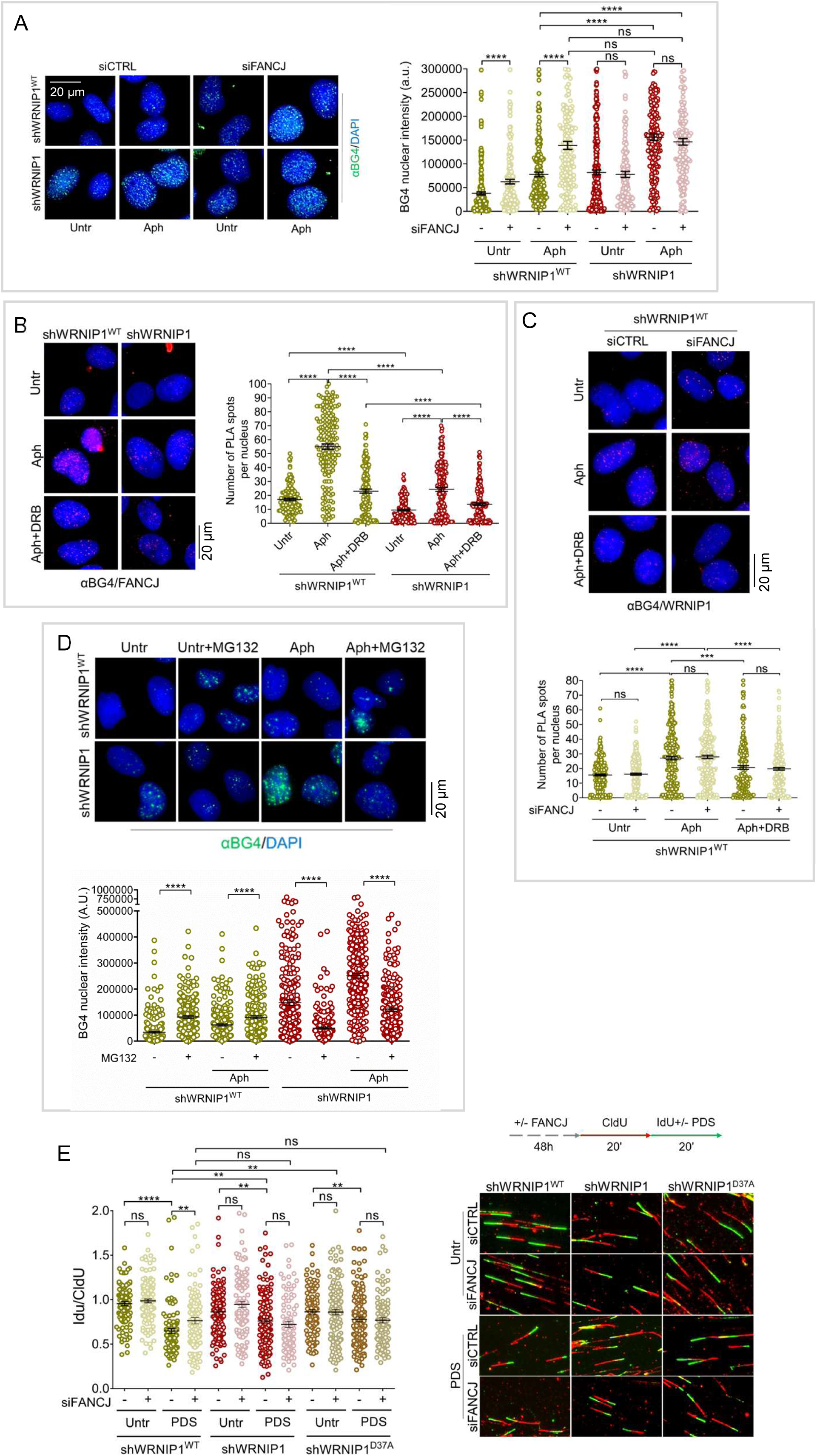
WRNIP1 and FANCJ cooperate to promote G4 resolution. (A) **G4 accumulation in FANCJ-depleted cells.** shWRNIP1^WT^ and shWRNIP1 cells were depleted of FANCJ by RNAi and treated or not with Aph for 24 h, fixed and stained with BG4 (G4s) antibody. DNA was counterstained with DAPI. Representative images are shown. Dot plot shows nuclear BG4 fluorescence intensity. Horizontal lines indicate median values (ns: *P* > 0.05; \*\*\*\**P* < 0.0001; Mann-Whitney test). (B) **G4-FANCJ interactions detected by PLA.** shWRNIP1^WT^ and shWRNIP1 cells were treated with or without Aph for 24 h and DRB during the last 3 h, fixed and subjected to PLA using BG4 (G4s) and anti-FANCJ antibodies. DNA was counterstained with DAPI. Representative images are shown. Dot plot shows the number of PLA spots per nucleus. Horizontal lines indicate median values (\*\*\*\**P* < 0.0001; Kruskal-Wallis test). (C) **G4-WRNIP1 interactions detected by PLA.** shWRNIP1^WT^ cells depleted of FANCJ by RNAi were treated as in (B). PLA was performed using BG4 (G4s) and anti-WRNIP1 antibodies. DNA was counterstained with DAPI. Representative images are shown. Dot plot shows the number of PLA spots per nucleus. Horizontal lines indicate median values (ns: *P* > 0.05; \*\*\**P* < 0.001; \*\*\*\**P* < 0.0001; Kruskal-Wallis test). (D) **G4 levels following proteasome inhibition.** shWRNIP1^WT^ and shWRNIP1 cells were treated or not with Aph for 24 h and MG132 during the last 6 h, fixed and stained with BG4 (G4s) antibody. DNA was counterstained with DAPI. Representative images are shown. Dot plot shows nuclear BG4 fluorescence intensity. Horizontal lines indicate median values (\*\*\*\**P* < 0.0001; Mann-Whitney test). (E) **Replication dynamics assessed by DNA fiber assay.** shWRNIP1^WT^, shWRNIP1 and shWRNIP1^D37A^ cells were sequentially pulse-labelled with CldU and IdU and treated or not with 5 μM PDS, as indicated. Representative DNA fiber images are shown. Dot plot shows the CldU/IdU ratios. Horizontal lines indicate median values (ns: *P* > 0.05; \*\**P* < 0.01; \*\*\*\**P* < 0.0001; Mann-Whitney test).

Mechanistically, WRNIP1 loss impaired FANCJ recruitment to G4 structures, as shown by reduced FANCJ-G4 PLA signal following Aph treatment or transcription inhibition (Figure 6B). This defect was rescued by proteasome inhibition (Supplementary Figure 7), indicating that WRNIP1 promotes FANCJ stability and/or retention at chromatin. In contrast, FANCJ depletion did not affect association of WRNIP1 with G4 as measured by PLA (Figure 6C), placing WRNIP1 upstream of FANCJ in this pathway. Importantly, stabilization of FANCJ reduced G4 accumulation in WRNIP1-deficient cells after Aph treatment (Figure 6D), further supporting this model.

Furthermore, replication fork analyses revealed that depletion of FANCJ, loss of WRNIP1 or mutation in the UBZ domain similarly reduced fork speed under G4-stabilizing conditions, with no additive effect upon co-depletion (Figure 6E).

Our findings indicate that WRNIP1 and FANCJ function in a common pathway that promotes G4 unwinding and supports replication fork progression. WRNIP1 appears to act upstream by stabilizing FANCJ and facilitating its association with chromatin; in its absence, impaired FANCJ function leads to G4 accumulation and replication stress.

### ATM protects FANCJ from ubiquitin-mediated degradation

FANCJ downregulation appears to result from proteasomal degradation, as treatment with MG132 restores its levels in WRNIP1-deficient and UBZ mutant cells. Since previous studies have shown that ATM can counteract ubiquitin-mediated protein degradation ^33,34^, we asked whether ATM contributes to FANCJ stabilization in this context.

Inhibition of ATM, but not ATR, markedly reduced FANCJ levels (Supplementary Figure 8A), indicating that ATM specifically supports FANCJ stability. Consistently, ATM inhibition also decreased FANCJ association with chromatin, as shown by fractionation assays, and this effect was reversed by MG132 (Figure 7A). Consistent results were obtained treating ATM-inhibited cells with PDS (Supplementary Figure 8B). These results suggest that, in the absence of ATM activity, FANCJ is targeted for proteasomal degradation, limiting its chromatin binding.

**Figure 7.**
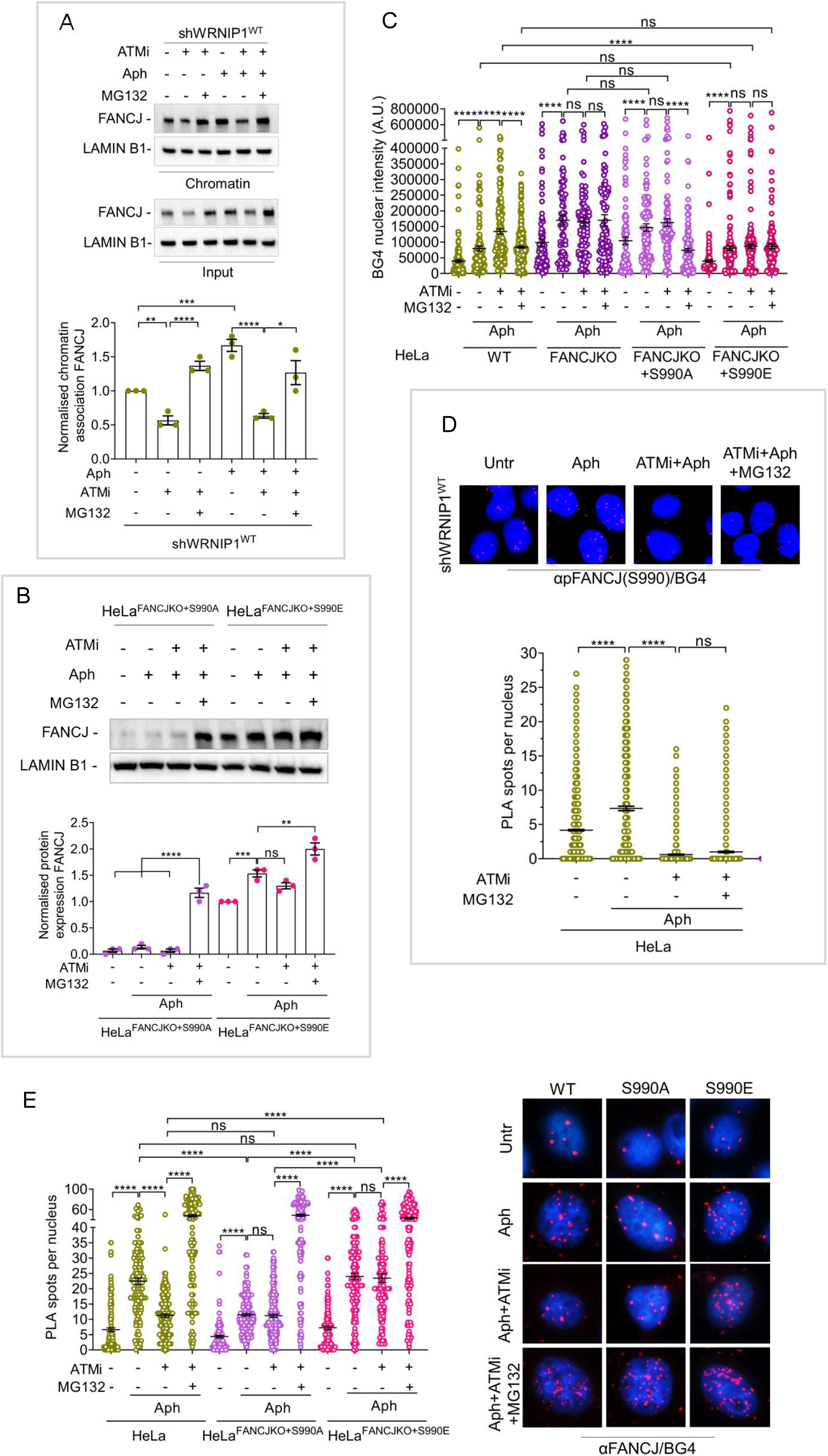
ATM-dependent stabilization of FANCJ (A) FANCJ association with chromatin following ATM inhibition. shWRNIP1^WT^ cells were pre-treated with 10 μM KU55933 (ATM inhibitor) for 1 h, treated or not with Aph for 24 h and 10 μM MG132 during the last 4 h. Chromatin fractions were immunoblotted with antibodies against FANCJ and LAMIN B1. Data are shown as super plot of FANCJ/LAMIN B1 ratios normalized to control (\**P* < 0.05; \*\**P* < 0.01; \*\*\**P* < 0.001; \*\*\*\**P* < 0.0001; one-way ANOVA test). **(B) Stability of mutant forms of FANCJ.** HeLa wild-type, HeLa FANCJ knockout (FANCJ^KO^) cells, FANCJ^KO^ cells complemented with a phosphorylation-deficient mutant (FANCJ^KO+S990A^) or a phosphomimetic mutant (FANCJ^KO+S990E^) were pre-treated with 10 μM KU55933 (ATM inhibitor) for 1 h, treated or not with 24 h Aph and 10 μM MG132 during the last 4 h. Blots were probed with antibodies against FANCJ and LAMIN B1. Data are shown as super plot of FANCJ/LAMIN B1 ratios normalized to control (ns: *P* > 0.05; \*\**P* < 0.01; \*\*\**P* < 0.001; \*\*\*\**P* < 0.0001; one-way ANOVA test). **(C) G4 levels following proteasome inhibition.** Cells were treated as in (B), fixed and stained with BG4 (G4s) antibody. Dot plot shows nuclear BG4 fluorescence intensity. Horizontal lines indicate median values (ns: *P* > 0.05; \*\*\*\**P* < 0.0001; Mann-Whitney test). **(D) G4-phospho-FANCJ Ser990 interactions detected by PLA.** HeLa cells were treated as in (B). PLA was performed using BG4 (G4s) and anti-phospho-Ser990 FANCJ antibodies. DNA was counterstained with DAPI. Representative images of are shown. Dot plot shows the number of PLA spots per nucleus. Horizontal lines indicate median values (ns: *P* > 0.05; \*\*\**P* < 0.001; Kruskal-Wallis test). **(E) G4-FANCJ interactions in FANCJ mutant cells detected by PLA.** HeLa, FANCJ^KO+S990A^ and FANCJ^KO+S990E^ cells were treated as in (B). PLA was performed using BG4 (G4s) and anti-FANCJ antibodies. DNA was counterstained with DAPI. Representative images are shown. Dot plot shows the number of PLA spots per nucleus. Horizontal lines indicate median values (ns: *P* > 0.05; \*\*\*\**P* < 0.0001; Kruskal-Wallis test).

To further support this model, we examined RAD18, a known regulator of ATM signaling ^35^. RAD18 depletion similarly reduced FANCJ chromatin loading under both basal and Aph-treated conditions, and this defect was rescued by MG132 (Supplementary Figure 8C and D). Moreover, RAD18 disruption increased Aph-induced DNA damage, which persisted in WRNIP1-deficient cells but not in wild-type cells (Supplementary Figure 8E). Notably, RAD18 depletion phenocopies WRNIP1 loss, supporting the idea that WRNIP1 functions within a RAD18-ATM-FANCJ signaling axis.

Impaired DNA damage resolution is likely due to defective G4 unwinding caused by reduced FANCJ function. In line with this, ATM inhibition decreased FANCJ proximity with G4s in wild-type cells (Supplementary Figures 8F). In contrast, WRNIP1-deficient and UBZ mutant cells, already compromised in ATM signaling and FANCJ stability, did not show further reduction upon ATM inhibition (Supplementary Figures 8F). Remarkably, MG132 treatment restored FANCJ-G4 interactions across all cell lines (Supplementary Figures 8F).

Therefore, our results demonstrate that the ATM pathway plays a critical role in maintaining FANCJ stability by protecting it from ubiquitination-mediated degradation, thereby ensuring its proper recruitment to chromatin and efficient resolution of G4 structures.

### ATM-mediated phosphorylation of FANCJ regulates its function

FANCJ is phosphorylated in response to various treatments ^36,37^ and bioinformatic analyses identify Ser990 as a potential phosphorylation site for multiple kinases, including ATM/CHK2. Given that ATM promotes FANCJ stability, we investigated whether phosphorylation at S990 regulates this effect.

HeLa FANCJ-deficient (FANCJ^KO^) cells were complemented with wild-type FANCJ, a phosphorylation-deficient mutant (S990A), or a phosphomimetic mutant (S990E). The S990A mutant showed reduced protein levels even under basal conditions, which were restored by proteasome inhibition (Figure 7B and Supplementary Figure 9A), indicating increased degradation. In contrast, S990E remained stable, although its levels decreased upon ATM inhibition, suggesting that ATM-dependent phosphorylation at S990 protects FANCJ from proteasomal turnover (Figure 7B).

Cycloheximide chase assays confirmed that S990A is intrinsically unstable and degrades more rapidly, whereas S990E displays stability comparable to wild-type FANCJ (Supplementary Figure 9B).

Functionally, S990A-expressing cells accumulated G4 structures to levels similar to FANCJ-deficient or ATM-inhibited cells, while MG132 treatment restored G4 resolution (Figure 7C). Consistently, PLA analysis using a pS990-FANCJ antibody showed reduced proximity between the S990A mutant and G4s, resembling ATM-inhibited wild-type cells, whereas the S990E mutant did not display this reduction and maintained a robust signal (Figure 7D). This defect was rescued by MG132, further linking FANCJ stability to its ability to deal with G4 structures. Western blot confirmed antibody specificity, showing reduced signal upon ATM inhibition, consistent with ATM-dependent phosphorylation of Ser990 (Supplementary Figure 9C). Accordingly, phosphorylated FANCJ showed proximity with G4s under basal conditions and increased upon Aph, but this was lost upon ATM inhibition (Figure 7E). Importantly, MG132-stabilized FANCJ under ATM inhibition was not phosphorylated, indicating selective degradation of the non-phosphorylated form (Figure 7E).

These phenotypes were mirrored by DNA damage (Supplementary Figure 9D). S990A cells showed levels comparable to FANCJ loss or ATM inhibition with no further effect of ATM inhibition, whereas S990E resembled wild-type. MG132 partially reduced DNA damage in wild-type and S990A cells.

Together, these results demonstrated that ATM-dependent phosphorylation of FANCJ at Ser990 prevents its degradation, enabling its accumulation at G4 structures and promoting efficient G4 resolution during replication stress.

### USP1 knockdown promotes the proteasomal degradation of FANCJ

USP1 is a deubiquitinase that removes mono- and polyubiquitin chains to regulate protein stability and function ^38^. Its depletion increases ubiquitination and promotes proteasomal degradation of target proteins. Since our data indicate that FANCJ is regulated through a proteasome-dependent mechanism, we investigated whether USP1 contributes to its stability.

USP1 knockdown reduced FANCJ levels in wild-type (siUSP1) cells under both unperturbed and Aph-treated conditions, phenocopying WRNIP1 deficiency (siCTRL); this effect was rescued by MG132 (Figure 8A). Notably, USP1 depletion did not further decrease FANCJ levels in WRNIP1-deficient (siUSP1) cells (Figure 8A).

**Figure 8.**
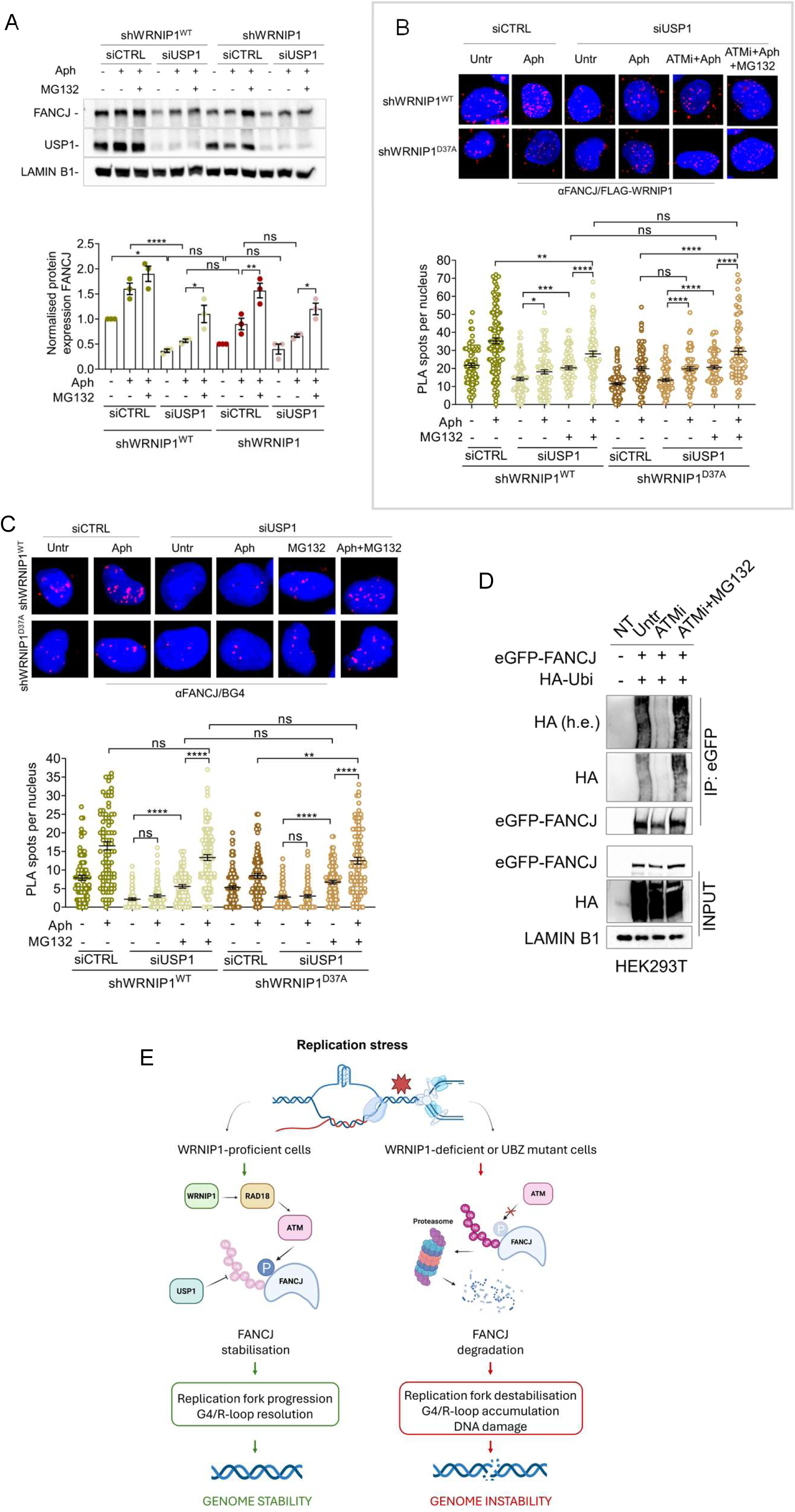
Proteasomal degradation of FANCJ (A) FANCJ stability in USP1-depleted cells. shWRNIP1^WT^and shWRNIP1 cells were depleted of USP1 by RNAi and, 48 h later, treated or not with Aph for 24 h, with 10 μM MG132 during the last 4 h. Blots were probed with antibodies against FANCJ, USP1 and LAMIN B1. Data are shown as super plot of FANCJ/LAMIN B1 ratios normalized to control (ns: *P* > 0.05; \**P* < 0.05; \*\**P* < 0.01; \*\*\*\**P* < 0.0001; one-way ANOVA test). **(B) WRNIP1-FANCJ interactions in USP1-depleted cells detected by PLA.** shWRNIP1^WT^ and shWRNIP1^D37A^ cells treated as in (A). PLA was performed using anti-FLAG (WRNIP1) and anti-FANCJ antibodies. DNA was counterstained with DAPI. Representative images are shown. Dot plot shows the number of PLA spots per nucleus. Horizontal lines indicate median values (ns: *P* > 0.05; \**P* < 0.05; \*\*\**P* < 0.001; \*\*\*\**P* < 0.0001; Kruskal-Wallis test). **(C) G4-FANCJ interactions in USP1-depleted cells detected by PLA.** Cells treated as in (A). PLA was performed using BG4 (G4s) and anti-FANCJ antibodies. DNA was counterstained with DAPI. Representative images are shown. Dot plot shows the number of PLA spots per nucleus. Horizontal lines indicate median values (ns: *P* > 0.05; \*\**P* < 0.01; \*\*\*\**P* < 0.0001; ANOVA test). **(D) Analysis of FANCJ ubiquitination.** HEK293T cells were co-transfected with control eGFP-FANCJ and HA-ubiquitin (HA-Ubi). At 48 h post-transfection, cells were treated with 20 μM KU55933 (ATM inhibitor) for 1 h, followed by 20 μM MG132 during the last 4 h. IP was performed with anti-eGFP antibody. Blots were probed with antibodies against Tag-HA, eGFP and LAMIN B1. **(E) WRNIP1 functions as a key regulator of FANCJ stability preventing genome instability.** Schematic representation of WRNIP1 function in replication stress response and genome stability maintenance. In WRNIP1-proficient cells, our data support a model in which the WRNIP1-RAD18-ATM axis ensures ATM-dependent FANCJ stabilisation, enabling efficient G4 resolution during replication stress, supporting replication fork progression, and preserving genome stability. Loss of WRNIP1 or mutation in its UBZ domain causes FANCJ destabilization and proteasomal degradation, resulting in G4 accumulation, impaired replication fork progression, DNA damage, and consequently genome instability.

We next examined FANCJ interactions by PLA. USP1 depletion reduced the PLA signal between FANCJ and WRNIP1 in wild-type cells to levels comparable to those observed in UBZ mutant cells, with no additional reduction upon USP1 knockdown in the mutant background (Figure 8B). MG132 restored this association in both contexts, supporting a role for USP1 in preventing FANCJ degradation. These results suggest that USP1 loss phenocopies the defect in ATM signaling observed in UBZ mutant cells.

Similarly, USP1 knockdown decreased the PLA signal between FANCJ and G4s in both wild-type and UBZ mutant cells, and this effect was reversed by MG132 (Figure 8C). While UBZ mutant cells already displayed reduced FANCJ-G4 interaction, USP1 depletion produced a stronger effect in wild-type cells, indicating that USP1 plays an important role in maintaining FANCJ association with G4 structures. The rescue by MG132 further supports the idea that proteasomal degradation is the primary mechanism limiting this interaction.

Furthermore, USP1 depletion promoted DNA damage accumulation, as shown by comet assay analysis (Supplementary Figure 10). Although USP1 loss did not further increase Aph-induced damage, stabilization of FANCJ by MG132 significantly reduced DNA damage in both cell lines, linking FANCJ stability to genome integrity.

Finally, we investigated whether ATM contributes to the regulation of FANCJ ubiquitination by performing immunoprecipitation assays using an HA-tagged ubiquitin plasmid ^39^. We found that FANCJ ubiquitination was already elevated under unperturbed conditions (Figure 8D). However, upon ATM inhibition, FANCJ protein levels decreased, likely due to enhanced protein degradation, and were restored following MG132 treatment. These findings suggest that ATM contributes to FANCJ stabilization by limiting its proteasome-dependent degradation.

Together, these findings identify USP1 as a regulator of FANCJ stability. By counteracting ubiquitination, USP1 prevents proteasomal degradation of FANCJ, thereby sustaining its interaction with WRNIP1 and G4 structures. Loss of USP1 compromises FANCJ function and contributes to genome instability under replication stress.

## DISCUSSION

G-quadruplexes (G4s) are non-canonical DNA structures that form throughout the genome and play regulatory functions. However, their persistence can compromise genome integrity by interfering with essential processes ^2,40,3^, promoting genomic instability and, in some contexts, tumorigenesis. Although several mechanisms involved in G4 resolution have been described, the pathways coordinating their timely removal during replication stress remain incompletely understood.

In this study, we identify a previously unrecognized role for WRNIP1 in promoting the stability and function of the G4 helicase FANCJ through activation of the ATM pathway. Our findings support a model in which ATM-dependent phosphorylation of FANCJ protects the helicase from ubiquitin-mediated proteasomal degradation, contributing to efficient resolution of G4 structures associated with R-loops.

Our data show that loss of WRNIP1 or mutation of its ubiquitin-binding zinc finger (UBZ) domain promotes the accumulation and persistence of R-loop-associated G4s, leading to transcription– replication conflicts (TRCs) and genome instability. In contrast, WRNIP1 ATPase activity is dispensable for G4 suppression and genome protection under these conditions, highlighting a specific role for the UBZ domain in G4 regulation.

Although the precise mechanism underlying G4 formation in this context remains unclear, R-loops are known to be enriched at genomic regions containing G4 motifs on the non-template strand ^41^. Consistent with this, we observe that G4s accumulate in close proximity to R-loops in a transcription-dependent manner under replication stress. One possible explanation is that G4 structures form within the displaced single-stranded DNA (ssDNA) generated by R-loops, as previously proposed ^10,42^. Supporting this model, PLA analyses indicate that G4 structures are closely associated with ssDNA-containing R-loop regions, suggesting that a substantial fraction of G4s accumulates at or near R-loops in WRNIP1-deficient and UBZ mutant cells.

The yeast homolog of WRNIP1, Mgs1, exhibits high affinity for G4 structures independently of its ATPase activity ^43^. Consistent with these observations, we find that WRNIP1 localizes in close proximity to G4s under replication stress conditions. Notably, this interaction does not require ATPase activity and is not significantly affected by mutation of the UBZ domain. In agreement with models linking G4 formation to impaired replication fork progression ^44^, we further show that G4 accumulation in WRNIP1-deficient or UBZ mutant cells occurs predominantly during S phase and depends on ongoing DNA synthesis, consistent with previous report ^26^.

How WRNIP1 prevents G4 accumulation at R-loops to support replication fork progression remains incompletely understood. WRNIP1 promotes restart of hydroxyurea-stalled replication forks through a mechanism requiring its ATPase activity ^19^. However, its role in limiting R-loop-associated TRCs is ATPase-independent and instead relies on the UBZ domain, which is also required for R-loop resolution ^21^. Since R-loop homeostasis is tightly regulated by ubiquitination ^45^ and WRNIP1 interacts with multiple ubiquitin signals ^46^, WRNIP1 may participate in a ubiquitination-dependent pathway that facilitates G4/R-loop resolution. Consistent with this possibility, WRNIP1 and its UBZ domain have been implicated in recruitment of proteins involved in checkpoint signalling and DNA repair, and ubiquitin binding is widely used by factors acting at stalled replication forks or during DNA repair ^47,48^. Together, these observations raise the possibility that WRNIP1 promotes G4 resolution by recruiting additional effector proteins through its ubiquitin-binding capacity. Importantly, although mutation of the UBZ domain does not impair WRNIP1 recruitment to G4s, it compromises downstream functionality.

G4 structures are unwound by specialized helicases ^49,50^. In yeast, the Pif1 helicase promotes recruitment of Mgs1 to G4s, although the molecular basis and functional significance of this cooperation remain incompletely understood ^43,51^. In mammalian cells, several G4-resolving helicases have been identified, including FANCJ, WRN, and BLM ^50,52,53,54^. Among these, FANCJ is considered one of the most efficient G4 helicases ^50^. Similar to WRNIP1, FANCJ suppresses G4 accumulation at replisomes ^55^ and promotes DNA synthesis across G4-containing regions in Xenopus egg extracts ^52^. Consistent with this, FANCJ deficiency results in G4 accumulation and DNA damage at G4-associated replication forks under mild replication stress ^55^, phenotypes that closely resemble those observed in WRNIP1-deficient and UBZ mutant cells. However, the mechanisms controlling FANCJ recruitment and stabilization at G4 sites remain poorly defined.

Unlike yeast Mgs1 and Pif1, WRNIP1 and FANCJ physically interact in mammalian cells under both basal and replication stress conditions, and this interaction is impaired by mutation of the WRNIP1 UBZ domain. Moreover, WRNIP1 deficiency markedly reduces the association of FANCJ with G4 structures, whereas WRNIP1-G4 association occurs independently of FANCJ. These findings suggest that WRNIP1 acts upstream of FANCJ at G4 sites and may facilitate FANCJ recruitment and/or stabilization. Consistent with this interpretation, chromatin-associated FANCJ levels are strongly reduced in the absence of WRNIP1 or its functional UBZ domain.

WRNIP1 has previously been described as an early responder to interstrand cross-links that promotes recruitment of the FANCD2/FANCI complex through direct protein interactions and UBZ-dependent functions ^47^(Socha et al., 2020). Similarly, WRNIP1 may facilitate FANCJ recruitment to G4 structures through its strong *in vitro* affinity for G4 DNA ^24^ and its ability to remain associated with G4-containing chromatin during replication stress, promoting efficient FANCJ function at these sites. Mechanistically, our findings support a model in which WRNIP1 and FANCJ operate within the same pathway to promote efficient G4 resolution. Loss of WRNIP1 or mutation of its UBZ domain compromises FANCJ chromatin association and stability, resulting in proteasome-dependent degradation of FANCJ and defective G4 unwinding. Consequently, G4 and R-loop structures persist following replication stress, delaying recovery and promoting genome instability. Consistent with this model, stabilization of FANCJ restores G4 resolution in WRNIP1-deficient cells. In contrast, combined disruption of WRNIP1-mediated ATM signalling and FANCJ does not further increase G4 accumulation, supporting epistasis within a common pathway.

Our findings further identify ATM as a regulator of FANCJ stability. ATM inhibition reduces both total and chromatin-associated FANCJ levels, and these effects are rescued by proteasome inhibition, suggesting that ATM signalling counteracts proteasomal turnover of FANCJ.

This observation is consistent with the broader role of ATM in coordinating DNA damage responses not only through checkpoint activation, but also through regulation of protein stability via post-translational modifications ^56^. ATM-dependent stabilization of DNA repair proteins has been described in multiple contexts, including regulation of p53 and replication-associated repair factors ^57,58^. More recently, similar mechanisms have been reported for additional genome maintenance proteins, supporting the idea that ATM-mediated phosphorylation can function as a protective signal against proteasomal degradation ^59,60,33,34^. The observation that proteasome inhibition restores FANCJ levels upon ATM inhibition is mechanistically informative, as it places ATM signalling upstream of the ubiquitin-proteasome system in regulating FANCJ turnover.

We further identify Ser990 as a critical ATM-dependent phosphorylation site controlling FANCJ stability and function. The non-phosphorylatable S990A mutant is unstable and undergoes proteasomal degradation, whereas the phosphomimetic S990E mutant is stabilized and functionally active. Previous studies have shown that FANCJ is phosphorylated in response to DNA damage and replication stress ^36,37^. Our results extend these findings by directly linking ATM-dependent phosphorylation to regulation of FANCJ stability.

Importantly, phosphorylation serves not only to protect FANCJ from degradation but also to promote its functional competence. Indeed, proteasome inhibition alone is insufficient to fully restore FANCJ activity in the absence of Ser990 phosphorylation, suggesting that this modification also contributes to productive chromatin engagement and/or protein-protein interactions ^61,62^.

We additionally identify USP1 as a deubiquitinase involved in maintaining FANCJ stability. USP1 depletion reduces FANCJ levels and impairs its association with both WRNIP1 and G4 structures, phenocopying WRNIP1 deficiency. USP1 is a major regulator of DNA damage tolerance pathways and controls ubiquitination of FANCD2 and PCNA ^38,63,64^. Our findings therefore expand the repertoire of USP1-regulated substrates and place USP1 within the WRNIP1-ATM regulatory axis controlling FANCJ homeostasis. This is consistent with the functional interplay between USP1 and the Fanconi anemia pathway, given that FANCJ is a bona fide Fanconi anemia gene product involved in interstrand cross-link repair ^65,66^.

The epistatic relationship observed between USP1 and WRNIP1 further supports the idea that these factors converge on regulation of FANCJ ubiquitination rather than functioning through independent parallel pathways. This is consistent with the broader interplay between ubiquitin signalling and ATM/ATR-dependent pathways in coordinating genome maintenance during replication stress ^67^ ^68^. Together, our data support a model in which the WRNIP1-RAD18-ATM axis cooperates with USP1 to maintain a stable chromatin-associated pool of FANCJ required for efficient G4 resolution during replication stress (Figure 8E).

Several questions remain to be addressed. Although our findings support a role for ATM-dependent phosphorylation in protecting FANCJ from proteasomal degradation, the E3 ubiquitin ligase(s) responsible for FANCJ turnover remain unknown. In addition, whether WRNIP1 directly recruits FANCJ to G4 structures or instead promotes a chromatin environment permissive for FANCJ loading will require further investigation.

Overall, our findings have potential clinical implications in cancer biology. Since replication stress and G4/R-loop accumulation are hallmarks of many tumor types, defects in the WRNIP1-ATM-FANCJ axis may contribute to tumor progression while simultaneously creating therapeutic vulnerabilities. In this context, tumors with impaired WRNIP1 pathway function may display increased sensitivity to G4-stabilizing compounds or replication stress-inducing therapies, opening potential opportunities for synthetic lethality.

## MATERIALS AND METHODS

### Cell lines and culture conditions

The SV40-transformed MRC5 fibroblast cell line (MRC5SV) stably expressing a WRNIP1-targeting shRNA (shWRNIP1), as well as isogenic cell lines stably expressing RNAi-resistant full-length wild-type WRNIP1 (shWRNIP1^WT^), the ATPase-dead mutant WRNIP1T294A (shWRNIP1^T294A^), and the ubiquitin-binding zinc finger (UBZ) domain mutant WRNIP1D37A (shWRNIP1^D37A^) were generated as previously described ^21^. Cells were cultured in the presence of neomycin (1 mg/mL) and puromycin (100 ng/mL) to maintain selective pressure for expression.

HEK293T and U2OS cell lines were obtained from the American Type Culture Collection (ATCC, VA, USA). HeLa cell lines that express wild-type FANCJ and the corresponding FANCJ knockout cells (HeLa KO) were kindly provided by Dana Branzei (IFOM ETS - The AIRC Institute of Molecular Oncology).

All cell lines were maintained in DMEM (Invitrogen) supplemented with 10% FBS (Boehringer, Mannheim), 100 U/mL of penicillin, and 100 μg/mL of streptomycin. Cells were cultured at 37°C and 5% CO₂. All cell lines were routinely tested for mycoplasma contamination and confirmed to be mycoplasma-free.

### Immunofluorescence

Immunofluorescence analysis of G-quadruplex structures (G4s) was performed as previously described ^26^, with minor modifications. Cells grown on glass coverslips were fixed with ice-cold 80% methanol in PBS for 15 min at -20°C and then washed twice with PBS. Cells were subsequently blocked with 10% FBS in PBS for 1 h and incubated overnight at 4°C with the anti-G-quadruplex antibody (BG4; Sigma-Aldrich; 1:200).

Immunostaining for RNA-DNA hybrids was performed as previously described ^21^. Briefly, cells were fixed in 100% methanol for 10 min at -20°C, washed three times with PBS, and pre-treated with RNase A (6 μg/mL) for 45 min at 37°C in 10 mM Tris-HCl (pH 7.5) supplemented with 0.5 M NaCl. Subsequently, cells were incubated for 90 min with RNase III (New England Biolabs; 1:150) in low-salt buffer (50 mM Tris-HCl pH 7.6, 75 mM KCl, 3 mM MgCl2, and 0.1% BSA), followed by blocking in 2% BSA/PBS overnight at 4°C. cells were incubated overnight at 4 °C with the anti-DNA-RNA hybrid antibody [S9.6] (Kerafast; 1:100).

Detection of parental ssDNA strand was performed by pre-labelling cells for 20 h with 100 µM IdU (Sigma-Aldrich), followed by washing in drug-free medium and treatment with 0.4 µM Aphidicolin for 24 h. Cells were then washed with PBS, permeabilized with 0.5% Triton X-100 for 10 min at 4°C, fixed with a 3% formaldehyde/2% sucrose solution for 10 min, and blocked with 3% BSA in PBS for 15 min, as previously described ^19^. Fixed cells were incubated with an anti-IdU antibody (mouse monoclonal anti-BrdU/IdU; clone B44, Becton Dickinson; 1:100).

Following primary antibody incubation, cells were washed twice with PBS and incubated with the appropriate secondary antibody: goat anti-mouse Alexa Fluor 488 or goat anti-rabbit Alexa Fluor 594 (Molecular Probes). Secondary antibody incubation was performed for 1 h at room temperature in a humidified chamber. DNA was counterstained with DAPI (0.5 μg/mL).

Images were randomly acquired using an Eclipse 80i Nikon fluorescence microscope equipped with a VideoConfocal (ViCo) system. For each experimental time point, at least 200 nuclei were analysed. Nuclear foci were scored at 40× magnification. Fluorescence intensity per nucleus was quantified using ImageJ. Parallel control samples incubated either with the appropriate normal serum or with secondary antibody alone confirmed that the observed fluorescence signals were not attributable to nonspecific staining or experimental artefacts.

### *In situ* PLA assay

In situ proximity ligation assay (PLA) was performed using the Duolink PLA kit (Sigma-Aldrich) according to the manufacturer’s instructions, and as previously described ^69^. Exponentially growing cells were seeded onto 8-well chamber slides (Nunc Lab-Tek) at a density of 1.5-2.5 × 10⁴ cells/well. Following the indicated treatments, cells were permeabilized with 0.5% Triton X-100 for 10 min at 4°C, fixed with 100% methanol for 10 min, and blocked with 3% BSA in PBS for 15 min. After washing with PBS, cells were incubated with the appropriate combination of primary antibodies. For R-loop detection by PLA using the anti-S9.6 antibody, cells were fixed with ice-cold methanol for 10 min and treated with RNase A (6 μg/mL) for 45 min at 37°C in 10 mM Tris-HCl (pH 7.5) supplemented with 0.5 M NaCl. Subsequently, cells were incubated with RNase III (New England Biolabs; 1:150) in low-salt buffer (50 mM Tris-HCl, pH 7.6, 75 mM KCl, 3 mM MgCl₂, and 0.1% BSA) for 90 min, followed by blocking with 2% BSA in PBS for 1 h at 37°C.

The primary antibodies used were: anti-FLAG (mouse monoclonal, Sigma-Aldrich; 1:250), anti-WRNIP1 (rabbit polyclonal, Bethyl; 1:500), anti-PCNA (rabbit polyclonal, Abcam; 1:500), anti-RNA polymerase II (mouse monoclonal, Santa Cruz Biotechnology; 1:200), recombinant anti-S9.6 (rabbit monoclonal, Kerafast; 1:200), anti-BG4 (mouse monoclonal, Merck-Millipore; 1:200), anti-IdU antibody (mouse monoclonal anti-BrdU/IdU; clone B44, Becton Dickinson; 1:100), anti-phospho-BRIP1 (FANCJ) Ser990 (rabbit polyclonal, Bioss; 1:500), and anti-FANCJ (rabbit polyclonal, Proteintech; 1:500). Negative controls were performed using only one primary antibody.

Samples were incubated with PLA secondary probes MINUS and PLUS (PLA Probe anti-Mouse PLUS and anti-Rabbit MINUS; Sigma-Aldrich). Incubation with antibodies and PLA probes was performed in a humidified chamber for 1 h at 37°C. The PLA probes were subsequently ligated using connector oligonucleotides to generate a template for rolling-circle amplification. During amplification, products were hybridized with a red fluorescently labelled oligonucleotide. Samples were mounted using ProLong Gold antifade reagent and nuclei were counterstained with DAPI (blue). Images were acquired randomly using an Eclipse 80i Nikon fluorescence microscope equipped with a Video Confocal (ViCo) system.

### Chromatin fractionation and immunoprecipitation

Chromatin fractionation and immunoprecipitation experiments were performed as previously described ^70^. Analysis of protein distribution within the chromatin fraction was carried out using a standard chromatin fractionation protocol ^71^. Cells (1.5 × 10^7^) were harvested by centrifugation (5 min, 1,300 × g, 4°C), and the pellets were washed twice with PBS (2 min, 1,300 × g, 4°C). Cell pellets were resuspended in buffer A (10 mM HEPES pH 7.9, 10 mM KCl, 1.5 mM MgCl₂, 0.34 M sucrose, 10% glycerol, 1 mM DTT, supplemented with protease inhibitor cocktail). Triton X-100 (0.1%) was added, and cells were incubated for 5 min on ice. Nuclei were collected by centrifugation (4 min, 1,300 × g, 4°C). The supernatant was discarded, and nuclei were washed once with buffer A before being lysed in buffer B (3 mM EDTA, 0.2 mM EGTA, 1 mM DTT, supplemented with protease inhibitor cocktail). Insoluble chromatin was collected by centrifugation (4 min, 1,700 × g, 4°C), washed once with buffer B, and centrifuged again under the same conditions. The final chromatin pellet was resuspended in 2× sample loading buffer (100 mM Tris/HCl pH 6.8, 100 mM DTT, 4% SDS, 0.2% bromophenol blue, and 20% glycerol), sonicated on ice, and boiled for 10 min at 95°C prior to western blot analysis.

For immunoprecipitation (IP) experiments, exponentially growing HEK293T cells were cultured overnight at a density of 2.5 × 10^6^ cells per 100 mm Petri dish and treated or left untreated as indicated. Following treatment, cells were collected by centrifugation, and cell pellets were resuspended in IP lysis buffer (0.5% Triton X-100, 150 mM NaCl, 1 mM EGTA, 50 mM Tris/HCl pH 8.0), freshly supplemented with protease and phosphatase inhibitor cocktails (SERVA), 15 U/mL benzonase (Sigma-Aldrich), and 1 mM MgCl₂. DNA was subsequently fragmented by passing the lysed suspension 5-10 times through a needle attached to a 2 mL syringe, followed by incubation for 40 min on ice. After centrifugation, lysates from each IP sample were incubated with 20 μL of anti-FLAG M2 magnetic beads (Sigma-Aldrich) overnight at 4°C. In parallel, 25 μL of ChromoTek GFP-Trap® Magnetic Agarose (Proteintech) were used according to the manufacturer’s instructions. The IP reactions were washed four times with IP buffer, incubated with 2× sample loading buffer (100 mM Tris/HCl pH 6.8, 100 mM DTT, 4% SDS, 0.2% bromophenol blue, and 20% glycerol) for 10 min at 95°C before western blot analysis.

### Statistical analysis

Statistical analysis was performed using Prism 8 (GraphPad Software). Details of the individual statistical tests are indicated in the figure legends. Statistical differences was assessed using the Mann-Whitney test, Kruskal-Wallis test, and one-way or two-way Anova test, as appropriate. In all cases, ns: not significant; *P* > 0.05; \**P* < 0.05; \*\**P* < 0.01; \*\*\**P* < 0.001; \*\*\*\**P* < 0.0001. All experiments were repeated at least three times unless otherwise indicated.

## Supporting information

main text plus figures

## Acknowledgments

This research was funded by Associazione Italiana per la Ricerca sul Cancro to A.F. (IG #30446) and to P.P. (IG #28972).

The authors express their gratitude to Dana Branzei (IFOM-FIRC Institute of Molecular Oncology), Nagaraju Ganesh (Indian Institute of Science), Edward Yeh (University of Texas) and R.J. Crouch (National Institutes of Health) for providing helpful reagents and advice.

To my brother Roberto, for his keen interest in scientific research.

## Author contributions

A.F. and P.P. conceived the experiments, wrote the manuscript, and acquired funding. A.F. designed and supervised the project. P.V. performed most of the experiments with the assistance of R.P. A.F. and P.P. contributed to the interpretation of the results and manuscript preparation. All authors reviewed and approved the final version of the manuscript.

