## Supplementary material for "WRNIP1-ATM signaling promotes G-quadruplex resolution by preserving FANCJ stability": main text plus figures

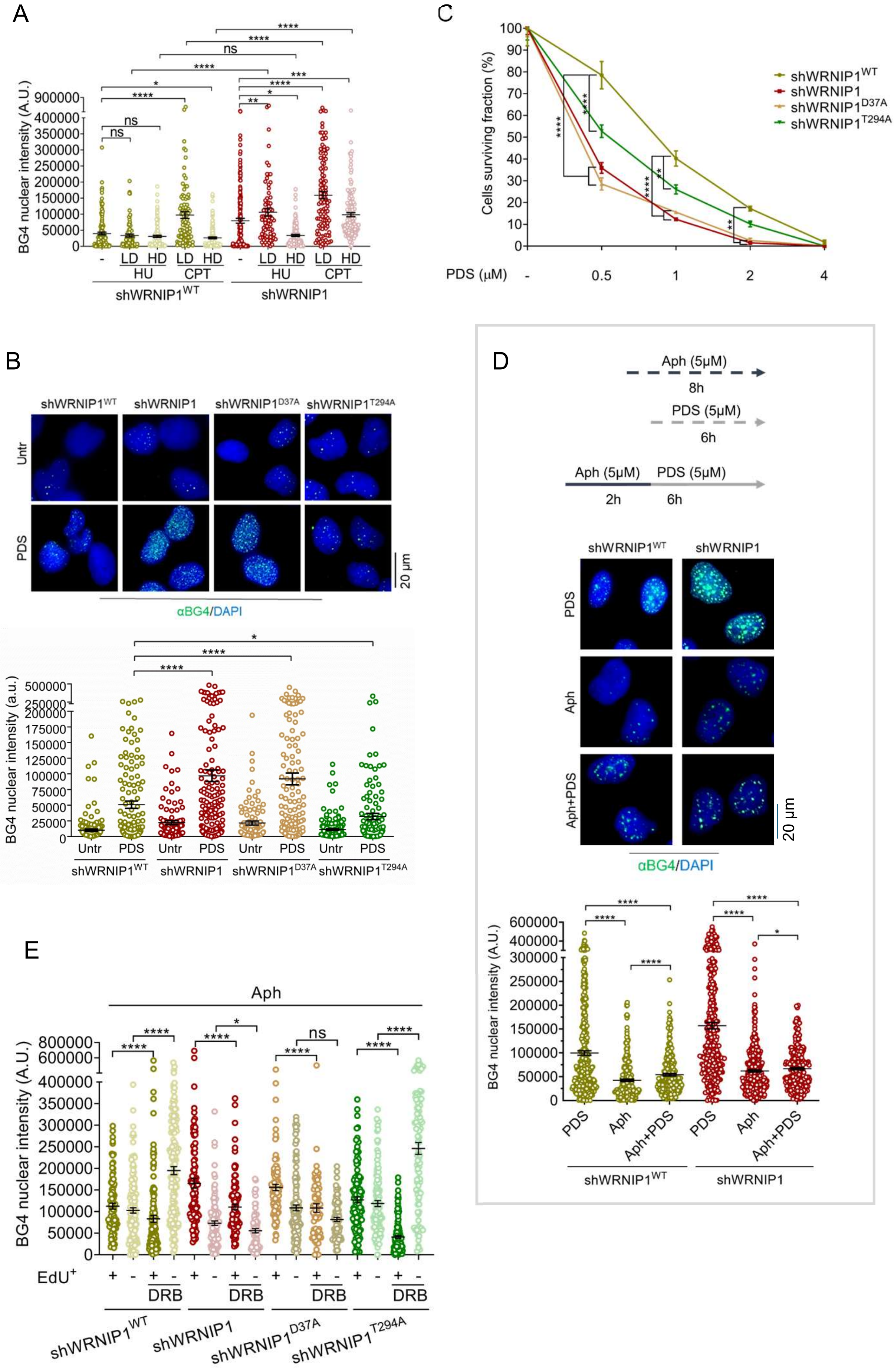


Suppl.Figure 1

A

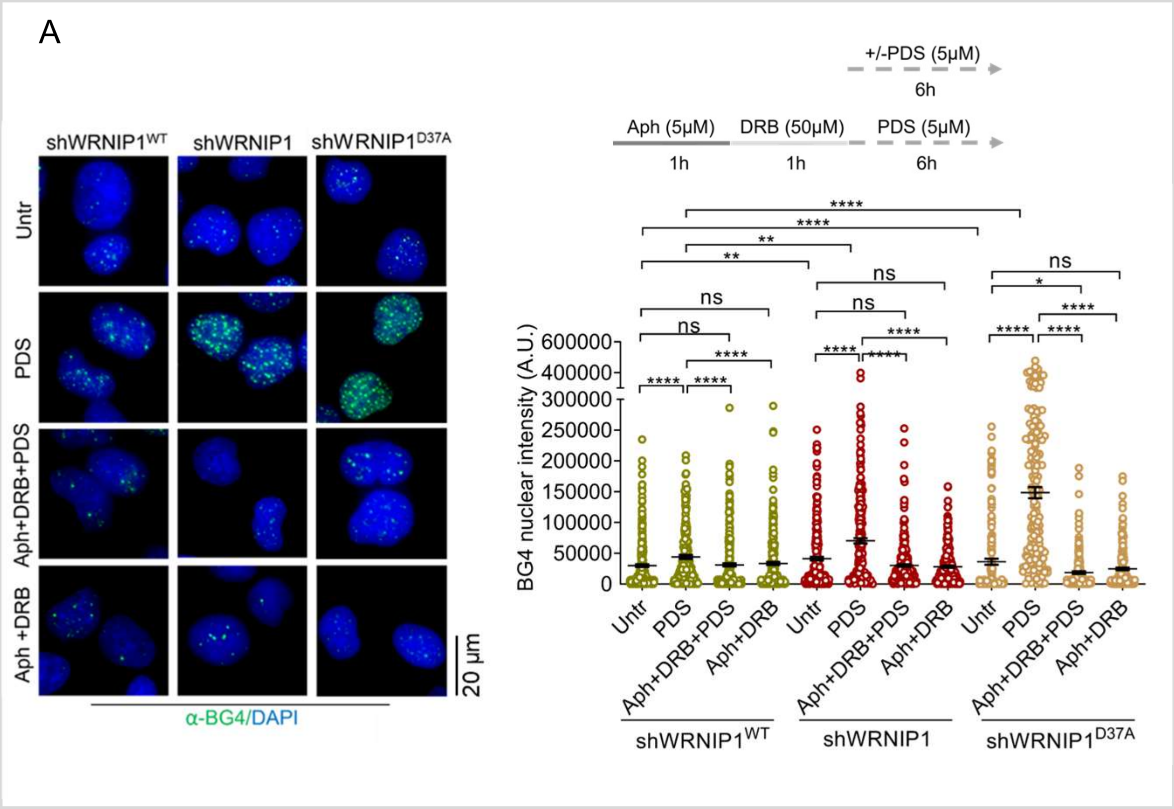


B

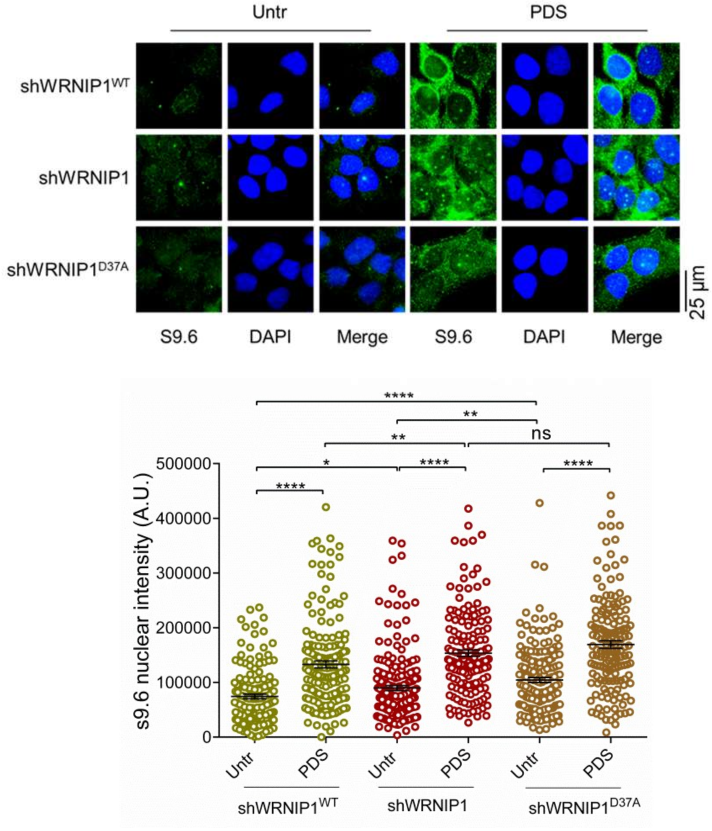


Suppl.Figure 2

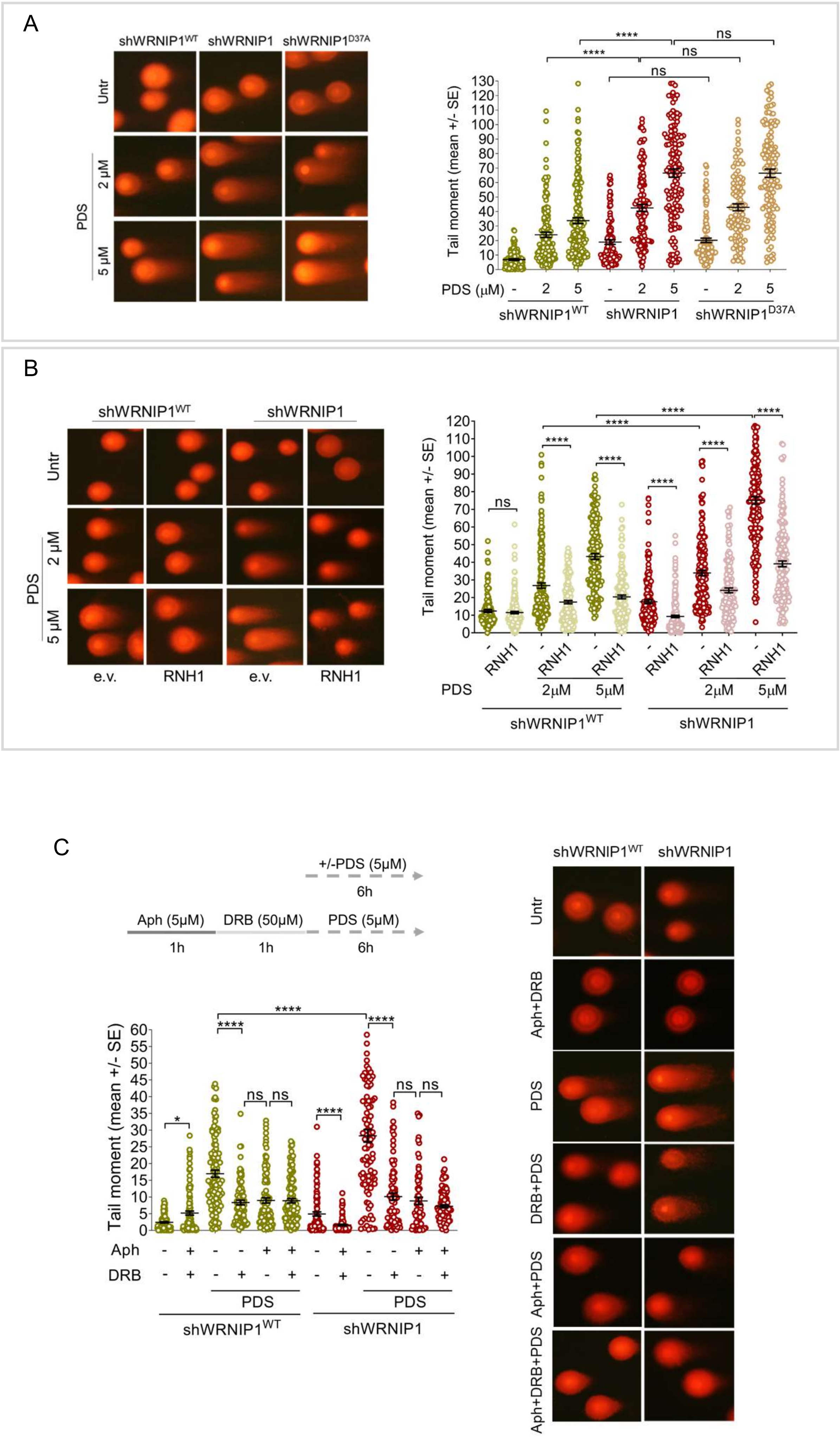


Suppl.Figure 3

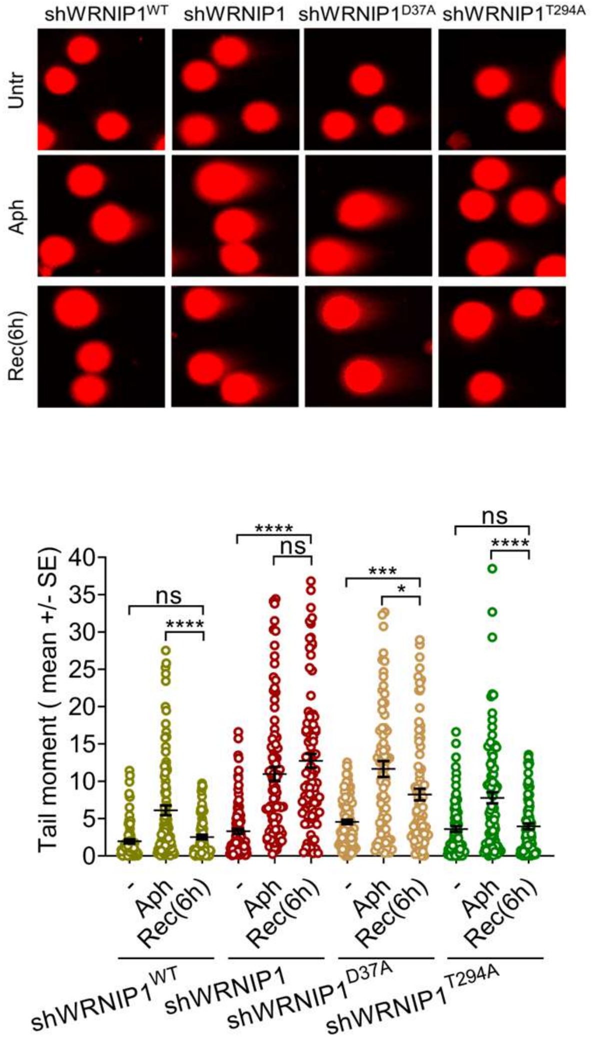


Suppl.Figure 4

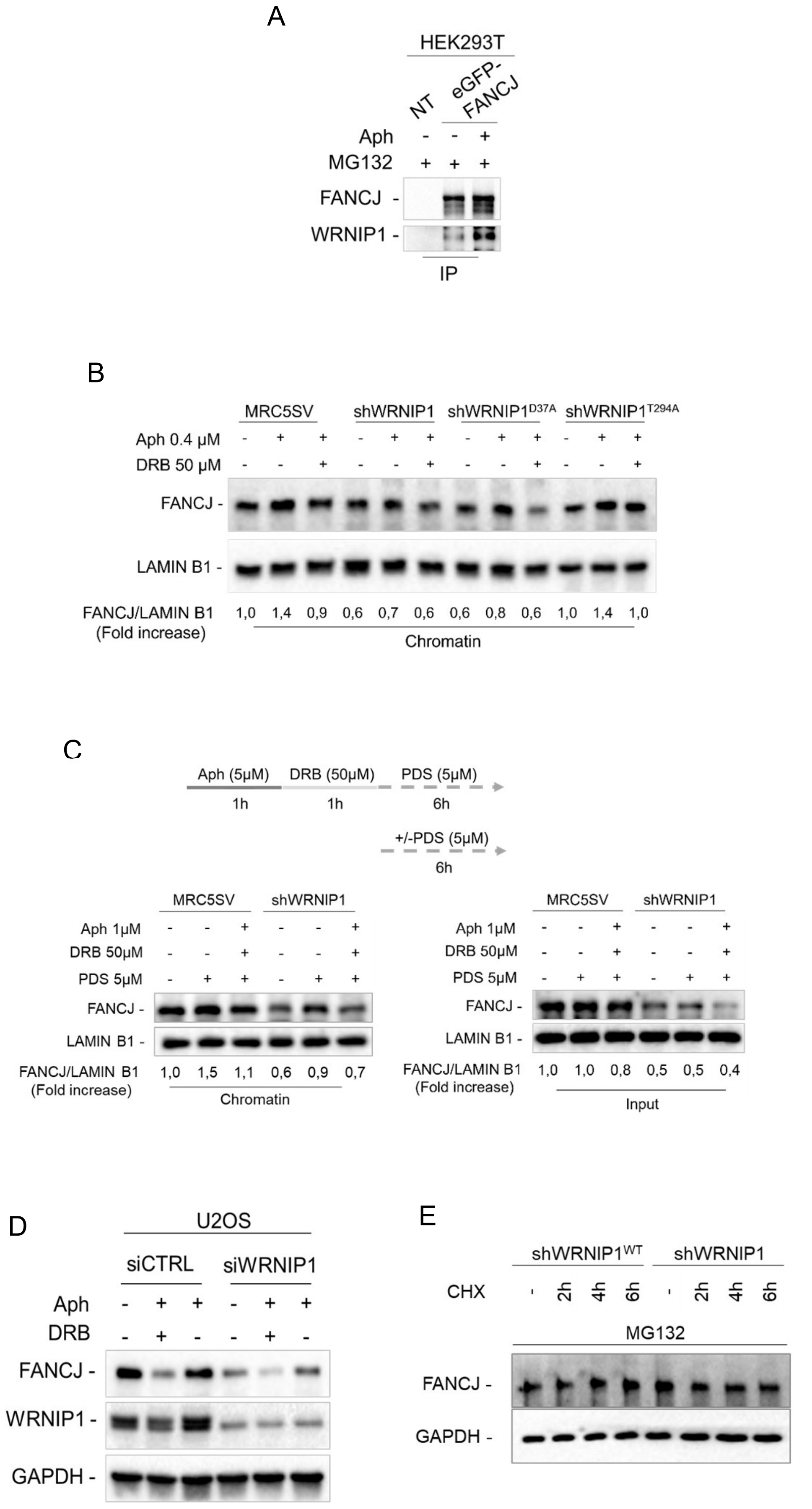


Suppl.Figure 5

A

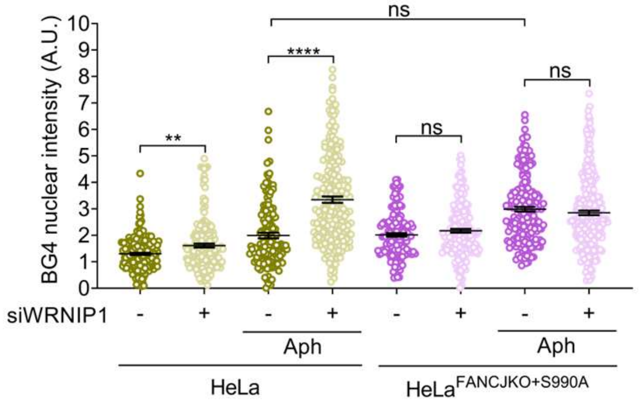


B

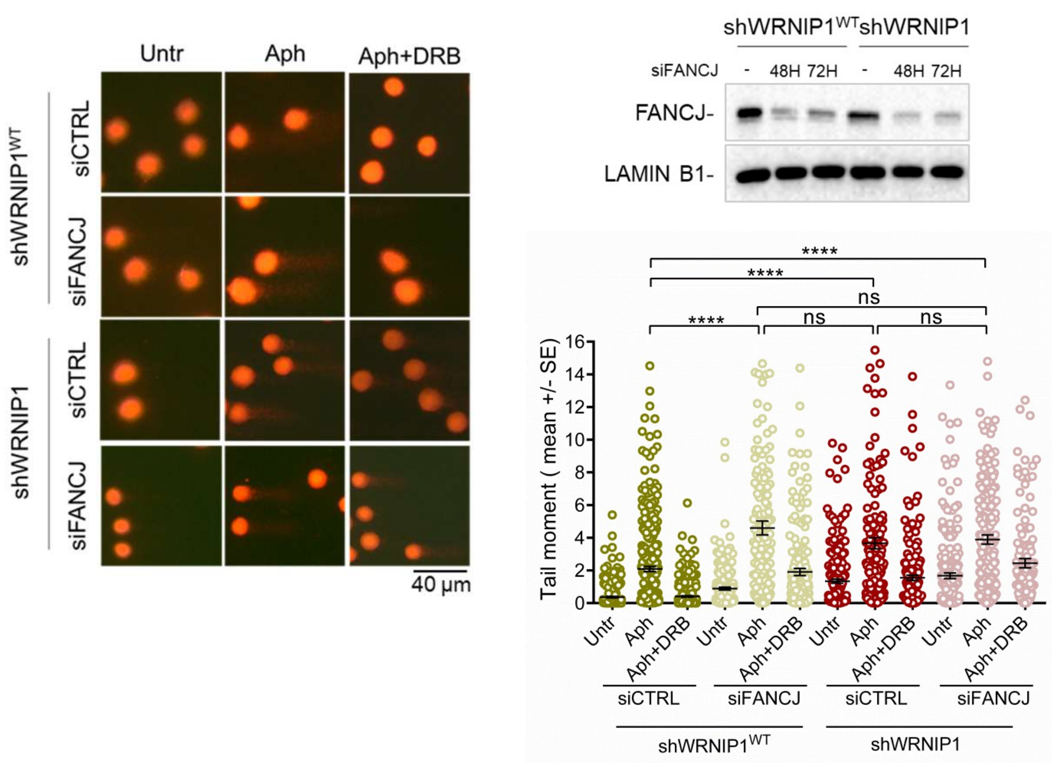


Suppl.Figure 6

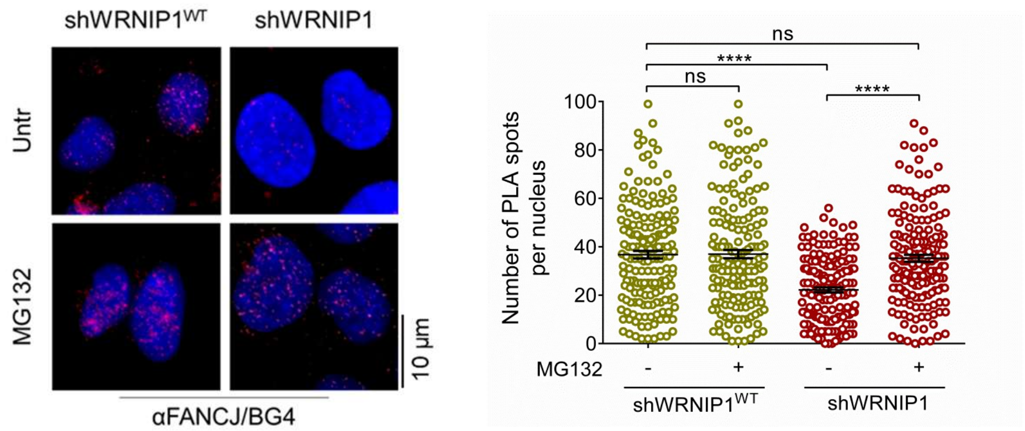


Suppl.Figure 7

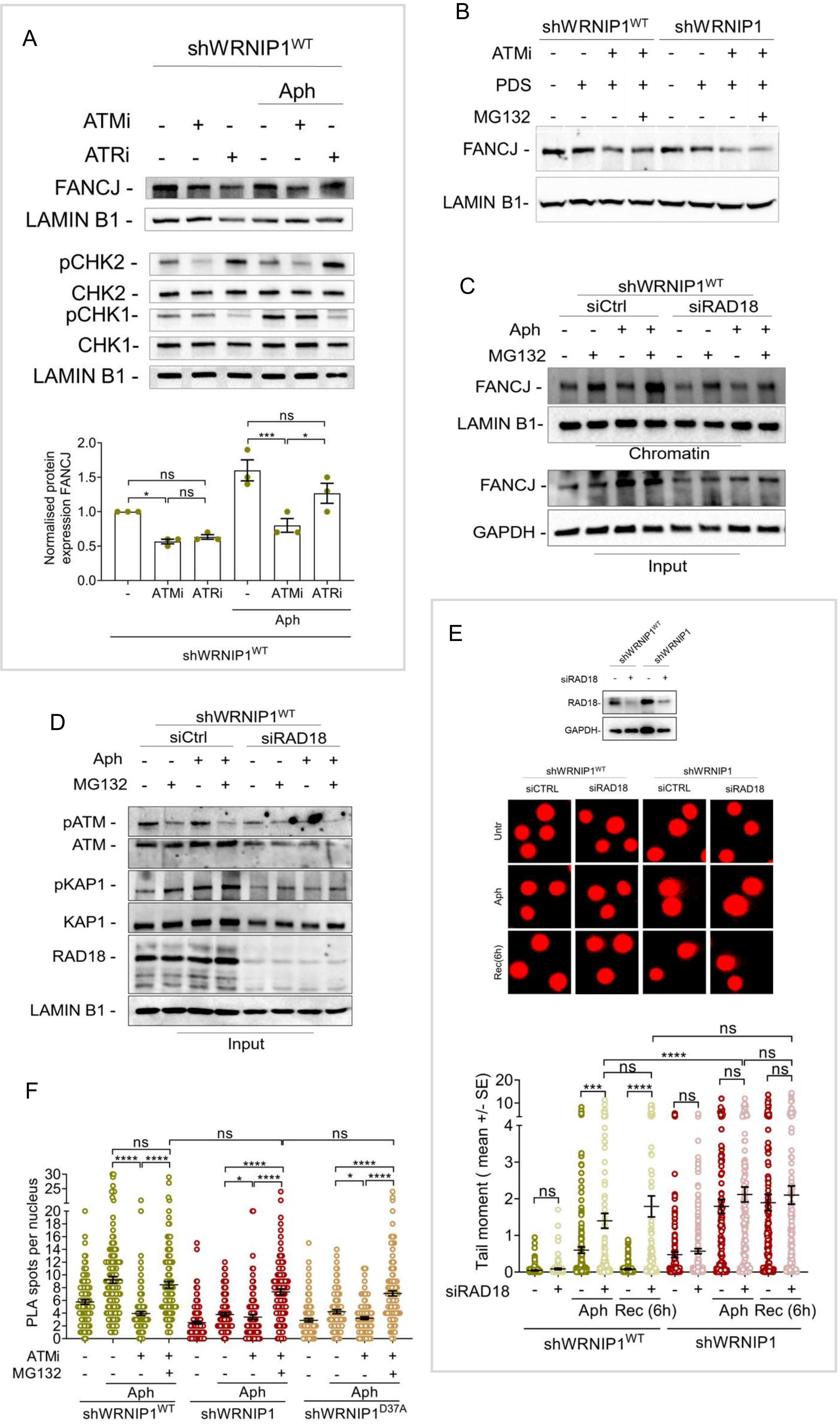


Suppl.Figure 8

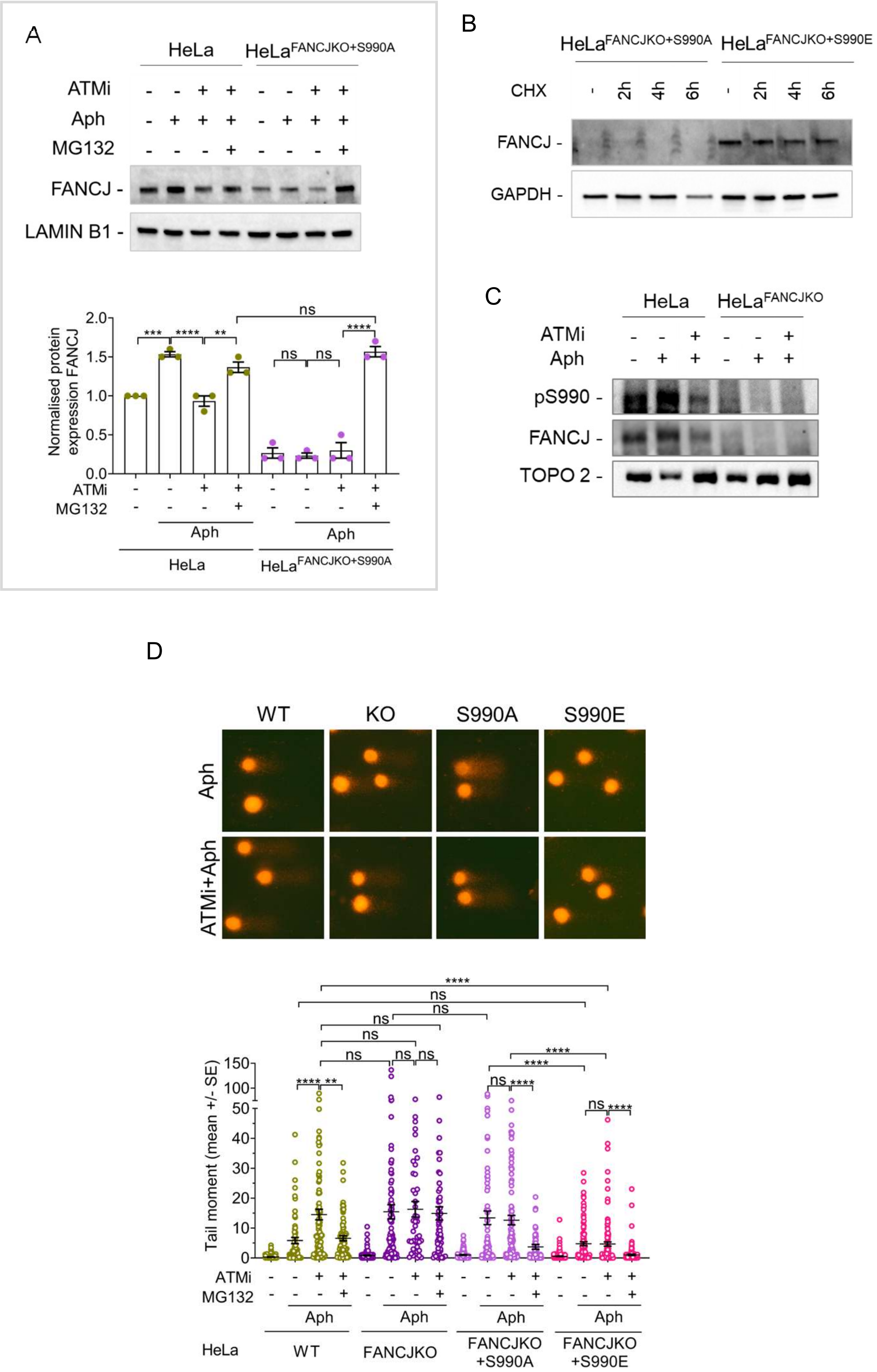


Suppl.Figure 9

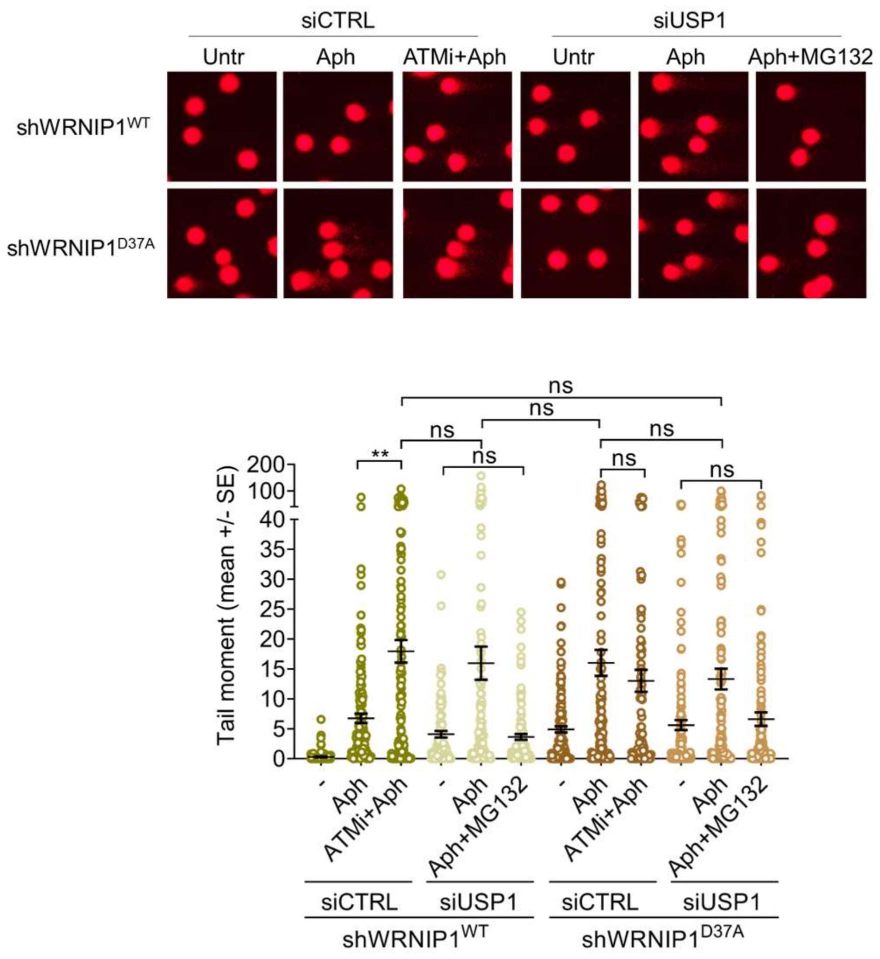


Suppl.Figure 10

### SUPPLEMENTARY MATERIALS AND METHODS

#### Chemicals

Chemicals were commercially obtained, and the final concentrations used were as follows: Aphidicolin (Aph; Sigma-Aldrich) at 0.4 and 5 µM; Hydroxyurea (HU; Sigma-Aldrich) at 0.5 and 4 mM; Camptothecin (CPT; Sigma-Aldrich) at 0.05 and 0.2 µM; 5,6-dichloro-1-β-D-ribofuranosylbenzimidazole (DRB; Sigma-Aldrich) at 50 µM; Pyridostatin trifluoroacetate salt (PDS; Selleck Chemicals) at 2 and 5 µM; MG132 (Selleck Chemicals) at 10 and 20 µM; 5-iodo-2′-deoxyuridine (IdU; Sigma-Aldrich) at 100 and 250 µM; 5-chloro-2′-deoxyuridine (CldU; Sigma-Aldrich) at 50 µM; ATM inhibitor KU-60019 (Sigma-Aldrich) at 10 and 20 µM; ATR inhibitor VE-821 (Selleck Chemicals) at 10 µM; and cycloheximide solution (CHX; Sigma-Aldrich) at 40 µg/mL. Stock solutions of all compounds were prepared in DMSO at concentrations above 1000× the final working concentration. The final concentration of DMSO in the culture medium was always maintained below 0.1%.

#### Plasmids and RNA interference

The construct used for RNaseH1 overexpression experiments was kindly provided by R.J. Crouch (National Institutes of Health). As previously described [1], the GFP-tagged RNaseH1 plasmid was generated by introducing a mutation at Met27 that abolishes the mitochondrial localization signal (RNaseH1-M27). The plasmid expressing FANCJ-S990E was kindly provided by Ganesh Nagaraju (Indian Institute of Science). pCDNA3.1 Bach1 (FANCJ) S990A was a gift from Sharon Cantor (Addgene plasmid #73643; http://n2t.net/addgene:73643; RRID:Addgene_73643). The pEGFP-C2-FANCJ plasmid was kindly provided by Yuliang Wu (University of Saskatchewan). The pCMV-FLAG-WRNIP1 plasmid (Open Biosystems) and the generation of the pCMV-FLAG-WRNIP1-D37A mutant plasmid has been previously described [2]. HA-Ubiquitin was a gift from Edward Yeh (Addgene plasmid #18712; http://n2t.net/addgene:18712; RRID:Addgene_18712). For plasmid expression, cells were transfected using the Neon™ Transfection System Kit (Invitrogen) according to the manufacturer's instructions for RNaseH1 expression, and using DreamFect (OZ Biosciences) for expression of HA-Ubi, eGFP-FANCJ, FANCJ-S990A, and FANCJ-S990E.

Genetic knockdown experiments targeting WRNIP1, FANCJ, USP1, and RAD18 were performed using Interferin (Polyplus) according to the manufacturer's instructions. siRNAs were used at the following final concentrations: 20 nM for WRNIP1, 10 nM for FANCJ, 20 nM for USP1, and 10 nM for RAD18.

WRNIP1 depletion was achieved using a QIAGEN siRNA targeting the 3′UTR region of the human transcript (5′-ATGAATTAATGTTATAAGG-3′). RAD18 knockdown was performed using a FlexiTube siRNA (QIAGEN) targeting the following mRNA sequence: 5′-ATGGTTGTTGCCCGAGGTTAA-3′. USP1 depletion was achieved by transfection with Silencer Select siRNA (Thermo Fisher Scientific; 5′-CCCTATGTATGAAGGATAT-3′). FANCJ knockdown was performed using ON-TARGETplus SMARTpool$^{TM}$ human BRIP1 siRNA (Horizon Discovery; L-010587-00-0005). As a negative control, a siRNA duplex targeting GFP was used.
Protein depletion efficiency was confirmed by western blotting using the appropriate antibodies.

**Western blotting**

Proteins were resolved by SDS-PAGE and transferred onto nitrocellulose membranes using the Trans-Blot Turbo Transfer System (Bio-Rad). Membranes were blocked with 5% non-fat dry milk (NFDM) in TBST (50 mM Tris/HCl pH 8.0, 150 mM NaCl, 0.1% Tween-20) and incubated with primary antibodies for 1 h at room temperature or overnight at 4°C for phospho-specific antibodies.
The following primary antibodies were used for western blotting: rabbit polyclonal anti-WRNIP1 (Bethyl Laboratories, 1:2500), mouse monoclonal anti-FLAG (Sigma-Aldrich, 1:1000), mouse monoclonal anti-GAPDH (Millipore, 1:5000), rabbit polyclonal anti-Lamin B1 (Abcam, 1:30,000), rabbit polyclonal anti-FANCJ (ProteinTech, 1:1500), mouse monoclonal anti-Cyclin A (Santa Cruz, 1:500), rabbit monoclonal anti-USP1 (Cell Signaling Technology, 1:1000), rabbit polyclonal anti-RAD18 (Abcam, 1:2000), rabbit polyclonal anti-phospho-BRIP1-Ser990 (Bioss, 1:2500), rabbit monoclonal anti-ATM (Cell Signaling Technology, 1:1000), mouse monoclonal anti-phospho-ATM-Ser1981 (Cell Signaling Technology, 1:1000), mouse monoclonal anti-eGFP (Santa Cruz, 1:500), rabbit monoclonal anti-phospho-CHK2-Thr68 (Cell Signaling Technology, 1:1000), mouse monoclonal anti-CHK2 (Calbiochem, 1:1000), rabbit polyclonal anti-phospho-KAP1-Ser824 (Bethyl Laboratories, 1:2000), rabbit polyclonal anti-KAP1 (Bethyl Laboratories, 1:2000), rabbit polyclonal anti-phospho-CHK1-Ser345 (Cell Signaling Technology, 1:1000), mouse monoclonal anti-CHK1 (Cell Signaling Technology, 1:1000), mouse monoclonal anti-HA (Thermo Fisher Scientific, 1:1500), and mouse monoclonal anti-Topoisomerase IIβ (Santa Cruz, 1:200).
The membranes were incubated with horseradish peroxidase (HRP)-conjugated goat species-specific secondary antibodies (Santa Cruz Biotechnology, 1:20,000) for 1 h at room temperature. Signal detection was performed using Western Bright ECL HRP substrate (Advansta), and images were acquired using a ChemiDoc XRS+ imaging system (Bio-Rad). Quantification was performed on scanned images of blots using ImageLab software (Bio-Rad). Quantifications are presented as mean

values from three independent experiments and normalized to the corresponding housekeeping proteins (LAMIN B1 or GAPDH).

#### Alkaline and neutral Comet assays

DNA damage induction was assessed by alkaline Comet assay (single-cell gel electrophoresis under denaturing conditions) as previously described [3]. Cellular DNA was stained with GelRed fluorescent dye (Biotium), and comets were visualized at 40× magnification using an Olympus fluorescence microscope. Slides were analysed using a computerized image analysis system (CometScore, Tritek Corp). The extent of DNA damage was quantified using the tail moment value, calculated as tail length × fraction of total DNA present in the tail. A minimum of 200 cells was analysed for each experimental condition. Apoptotic cells, characterized by a reduced comet head size and an extremely enlarged comet tail, were excluded from the analysis to avoid artificial overestimation of DNA damage.

The induction of DNA double-strand breaks (DSBs) was evaluated by neutral Comet assay as previously described [4]. Cellular DNA was stained with GelRed fluorescent dye (Biotium), and slides were analysed using the same imaging and quantification procedures described above.

#### DNA fiber assay

Cells were pulse-labelled with 50 μM 5-chloro-2′-deoxyuridine (CldU) and 250 μM 5-iodo-2′-deoxyuridine (IdU) at the indicated time points, with or without treatments as specified in the experimental schemes. DNA fibres were prepared and spread as previously described [2]. For immunodetection of labelled DNA tracks, the following primary antibodies were used: anti-CldU (rat monoclonal anti-BrdU/CldU; BU1/75 ICR1, Abcam; 1:100) and anti-IdU (mouse monoclonal anti-BrdU/IdU; clone B44, Becton Dickinson; 1:10). Secondary antibodies used were goat anti-mouse Alexa Fluor 488 and goat anti-rabbit Alexa Fluor 594 (Molecular Probes; 1:200). Antibody incubations were performed in a humidified chamber for 1 h at room temperature.

Images were randomly acquired from fields containing untangled DNA fibres using an Eclipse 80i Nikon fluorescence microscope equipped with a Video Confocal (ViCo) system. The lengths of red- and green-labelled tracks were measured using ImageJ software. Fork protection efficiency was calculated using the IdU/CldU ratio, where a ratio close to 1 indicates efficient fork protection and a ratio below 1 indicates fork degradation (Bhattacharya and Ganesh, 2026). A minimum of 100 individual fibres were analysed for each experiment, and the mean values from at least three independent experiments are presented. Statistical analyses were performed using GraphPad Prism software.

**Cell survival**

Clonogenic assays were performed by seeding 1,000 cells in 60-mm Petri dishes. The following day, cells were treated with different concentrations of PDS for 24 h, washed, and allowed to grow for an additional 2 weeks. Colonies were then fixed with methanol: acetic acid (3:1, v/v) and stained with 5 mg/mL Giemsa solution (Sigma). Colony counting was performed using a stereomicroscope. Only colonies containing more than 50 cells were considered for quantification. Cell survival was calculated relative to untreated control cells.

**Click-iT Cell Reaction (EdU detection)**

Cells were incubated with 125 μM EdU (5-ethynyl-2′-deoxyuridine) for 8 min before fixing to label newly synthesized DNA. Where indicated, cells were treated with Aph or 5,6-dichlorobenzimidazole riboside (DRB) to inhibit DNA replication or transcription, respectively, prior to and during EdU incorporation.

Following treatment, cells were washed with PBS and fixed with 4% paraformaldehyde for 10-15 min at room temperature. Cells were then permeabilized using 0.2-0.5% Triton X-100 in PBS for 10-15 min. The Click-iT reaction was performed according to the manufacturer's instructions. Briefly, cells were incubated with the Click-iT reaction cocktail containing $CuSO_4$, fluorescent azide dye, ascorbic acid, and reaction buffer additive, allowing covalent conjugation of the fluorescent probe to incorporated EdU via copper-catalyzed azide-alkyne cycloaddition. After incubation, cells were washed extensively with PBS and counterstained with DAPI. Samples were mounted using antifade mounting medium and imaged by fluorescence microscopy. Signal intensity or number of positive nuclei was quantified using image analysis software.

### SUPPLEMENTARY LEGENDS TO FIGURES

**Figure S1. WRNIP1 loss or mutation of its UBZ domain results in G4 accumulation**

**(A) G4 levels in HU- and CPT-treated cells at low and high doses.** shWRNIP1$^{WT}$ and shWRNIP1 cells were treated for 4 h with low (LD) or high dose (HD) of Hydroxyurea (HU) or Camptothecin (CPT) (0.5 and 4 mM HU; 0.05 and 0.2 µM CPT, respectively). Cells were fixed and stained with the BG4 (G4s) antibody. Dot plot shows nuclear BG4 fluorescence intensity. Horizontal lines indicate the median values (ns: $P > 0.05$; $*P < 0.05$; $**P < 0.01$; $****P < 0.0001$; Mann-Whitney test).

**(B) Analysis of G4 levels following PDS treatment.** shWRNIP1$^{WT}$, shWRNIP1, shWRNIP1$^{D37A}$ and shWRNIP1$^{T294A}$ cells were treated or not with 5 µM PDS for 24 h, fixed and stained with BG4 (G4s) antibody. DNA was counterstained with DAPI. Representative images are shown. Dot plot shows nuclear BG4 fluorescence intensity. Horizontal lines indicate the median values ($*P < 0.05$; $****P < 0.0001$; Mann-Whitney test).

(**C**) **Analysis of cell survival.** Cells were treated with increasing concentrations of PDS for 24 h, then washed and allowed to grow for an additional 2 weeks. Graph shows cell survival fractions ($**P < 0.01$; $****P < 0.0001$; two-way Anova test).

(**D**) **Effect of replication inhibition on G4 accumulation.** shWRNIP1$^{WT}$ and shWRNIP1 cells were treated with PDS, Aph, or both sequentially (Aph+PDS), as indicated. Cells were fixed and stained with BG4 (G4s) antibody. DNA was counterstained with DAPI. Representative images are shown. Dot plot shows nuclear BG4 fluorescence intensity. Horizontal lines indicate the median values ($*P < 0.05$; $****P < 0.0001$; Mann-Whitney test).

(**E**) **G4 formation predominantly occurs during S phase.** Cells were labelled with 10µM EdU for 30 min before fixation and treated or not with 0.4 µM Aph for 24 h, with DRB during the last 3 h. Cells were then fixed and subjected to Click-iT Cell Reaction (EdU detection). Dot plot shows nuclear BG4 fluorescence intensity in EdU positive (EdU+) and EdU negative (-) cells. Horizontal lines indicate the median values (ns: $P > 0.05$; $*P < 0.05$; $****P < 0.0001$; Mann-Whitney test).

**Figure S2**. **PDS increases G4 and R-loop accumulation in the absence of WRNIP1**

(**A**) **Impact of replication and transcription inhibition on G4 accumulation**. shWRNIP1$^{WT}$, shWRNIP1 and shWRNIP1$^{D37A}$ cells were treated as described in the scheme. Cells were fixed and stained with BG4 (G4s) antibody. DNA was counterstained with DAPI. Representative images are shown. Dot plot shows nuclear BG4 fluorescence intensity. Horizontal lines indicate median values (ns: $P > 0.05$; $*P < 0.1$; $**P < 0.01$; $****P < 0.0001$; Mann-Whitney test).

(**B**) **Evaluation of R-loop accumulation upon short PDS treatment.** shWRNIP1$^{WT}$, shWRNIP1 and shWRNIP1$^{D37A}$ cells were treated or not with 5 μM PDS for 6 h, then fixed and stained with the S9.6 antibody to detect R-loops. DNA was counterstained with DAPI. Representative images are shown. Dot plot shows nuclear S9.6 fluorescence intensity. Horizontal lines indicate median values (ns: $P > 0.05$; **$P < 0.01$; ****$P < 0.0001$; Mann-Whitney test).

**Figure S3**. **PDS-induced DSB formation depends on R-loops and replication/transcription**

(**A**) **PDS-induced double-strand breaks (DSBs) evaluated by neutral Comet assay.** shWRNIP1$^{WT}$, shWRNIP1 and shWRNIP1$^{D37A}$ cells were treated or not with 2 or 5 μM PDS for 24 h. Representative images are shown. Dot plot shows tail moment values. Horizontal lines indicate median values (ns: $P > 0.05$; ****$P < 0.0001$; Mann-Whitney test).

(**B**) **Rescue of PDS-induced DSBs by RNaseH1 overexpression by neutral Comet assay.** shWRNIP1$^{WT}$ and shWRNIP1 cells were transfected with GFP-tagged RNaseH1 (RNH1) or empty vector (e.v.) and treated or not with 2 or 5 μM PDS for 24 h. Representative images are shown. Dot plot shows tail moment values. Horizontal lines indicate median values (ns: $P > 0.05$; ****$P < 0.0001$; Mann-Whitney test).

(**C**) **Effect of replication and transcription inhibition on PDS-induced DSB formation.** shWRNIP1$^{WT}$ and shWRNIP1 cells were treated as described in the scheme, and subjected to neutral Comet assay. Representative images are shown. Dot plot shows tail moment values. Horizontal black lines represent the median values (ns: $P > 0.05$; *$P < 0.05$; ****$P < 0.0001$; Mann-Whitney test).

**Figure S4. WRNIP1-deficient or UBZ mutant cells fail to recover from Aph-treatment**

**Detection of DSBs by neutral Comet assay.** shWRNIP1$^{WT}$, shWRNIP1 and shWRNIP1$^{D37A}$ cells were treated or not with Aph for 24 h and allowed to recover in drug-free medium for 6 h. Representative images are shown. Dot plot shows tail moment values. Horizontal lines indicate median values (ns: $P > 0.05$; *$P < 0.05$; ***$P < 0.001$; ****$P < 0.0001$; Mann-Whitney test).

**Figure S5**. **Loss of WRNIP1 or mutation of its UBZ domain reduces FANCJ levels**

**(A) Co-immunoprecipitation of FANCJ and WRNIP1.** HEK293T cells were transfected with eGFP-FANCJ and treated or not with 0.4 μM Aph for 24 h and 20 μM MG132 during the last 4 h. IP was performed using an anti-eGFP Nanotrap beads. Blots were probed with antibodies against WRNIP1 and FANCJ.

(**B**) **Chromatin-bound FANCJ upon transcription inhibition.** MRC5SV, shWRNIP1, shWRNIP1$^{D37A}$ and shWRNIP1$^{T294A}$ cells were treated or not with Aph and DRB during the last 3 h. Chromatin fractions were immunoblotted with antibodies against FANCJ and LAMIN B1. Chromatin-bound FANCJ levels are expressed as the FANCJ/LAMIN B1 ratios, normalized to the untreated control.

(**C**) **FANCJ association with chromatin following combined replication and transcription inhibition.** MRC5SV and shWRNIP1 cells were treated as indicated in the scheme. Chromatin fractionation and immunoblotting were performed as in (B).

(**D**) **FANCJ levels in WRNIP1-depleted U2OS cells.** U2OS cells were depleted of WRNIP1 by RNAi and, 48 h hours later, treated or not with Aph for 24 h and DRB during the last 3 h. Blots were probed with antibodies against FANCJ, WRNIP1 and GAPDH. WRNIP1 knockdown efficiency was confirmed by Western blotting. FANCJ and WRNIP1 levels are expressed as the FANCJ/GAPDH and WRNIP1/GAPDH ratios, respectively, normalized to the untreated control.

**(E) Analysis of FANCJ stability**. shWRNIP1$^{WT}$ and shWRNIP1 cells were treated or not with 40 μg/mL cycloheximide and 10μM MG132 as indicated to assess FANCJ stability. Blots were probed with antibodies against FANCJ and GAPDH.

**Figure S6. FANCJ depletion exacerbates DNA damage accumulation upon replication stress**

(**A**) **G4 levels in WRNIP1-depleted cells expressing a phosphorylation-deficient FANCJ mutant.** HeLa and FANCJ$^{KO+S990A}$ cells were depleted of WRNIP1 by RNAi and, 48 h later, treated or not with Aph for 24 h, fixed and stained with BG4 (G4s) antibody. Dot plot shows nuclear BG4 fluorescence intensity. Horizontal lines indicate the median values (ns: $P > 0.05$; $^{**}P < 0.01$; $^{****}P < 0.0001$; Mann-Whitney test).

(**B**) **Evaluation of Aph-induced DNA damage in FANCJ-depleted cells by alkaline Comet assay.** FANCJ-depleted cells were treated or not with Aph for 24 h, with DRB during the last 6 h. FANCJ knockdown efficiency was confirmed by Western blotting. Blots were probed with antibodies against FANCJ and LAMIN B1. Representative images are shown. Dot plot shows tail moment values. Horizontal lines indicate median values (ns: $P > 0.05$; $^{****}P < 0.0001$; Mann-Whitney test).

**Figure S7. Proteasome inhibition enhances FANCJ-G4 co-localization of upon replication stress**

**G4-FANCJ interactions following proteasome inhibition detected by PLA.** shWRNIP1$^{WT}$ and shWRNIP1 cells were treated or not with Aph for 24 h, with MG132 added during the last 4 h. PLA was performed using BG4 (G4s) and anti-FANCJ antibodies. DNA was counterstained with DAPI.

Representative images are shown. Dot plot shows the number of PLA spots per nucleus. Horizontal lines indicate median values (ns: $P > 0.05$; ***$P < 0.001$; ****$P < 0.0001$; Kruskal-Wallis test).

**Figure S8. ATM signaling regulates FANCJ function and DNA damage responses**

(A) **Effect of ATM or ATR inhibition on FANCJ levels.** Cells were pre-treated with 10 µM KU55933 (ATM inhibitor) or 10 µM VE-821 (ATR inhibitor) for 1 h and treated or not with 24 h Aph. Blots were immunoblotted for indicated antibodies. FANCJ/LAMIN B1 ratios were normalized to untreated controls and quantified (ns: $P > 0.05$; *$P < 0.05$; ***$P < 0.001$; one-way ANOVA test).

**(B) Effect of ATM inhibition on FANCJ levels following PDS treatment.** Cells were pre-treated with 10 µM KU55933 (ATM inhibitor) for 1 h, treated or not with 2µM PDS for 24 h and 10 µM MG132 during the last 4 h. Blots were immunoblotted for FANCJ and LAMIN B1.

**(C) Requirement of RAD18 for FANCJ chromatin recruitment.** RAD18-depleted shWRNIP1$^{WT}$ cells were treated or not with Aph for 24 h, and with 10 µM MG132 during the last 4 h. Chromatin fractions were immunoblotted for indicated antibodies. Chromatin-bound FANCJ was quantified as FANCJ/LAMIN B1 relative to untreated cells.

(**D**) **ATM pathway activation in RAD18-depleted cells.** RAD18-depleted shWRNIP1$^{WT}$ cells were treated or not with Aph for 24 h, with 10 µM MG132 during the last 4 h. Blots immunoblotted for indicated antibodies. RAD18 knockdown efficiency was confirmed by Western blotting.

**(E) Recovery from Aph-induced DNA damage in RAD18-depleted cells**. RAD18-depleted cells were treated or not with Aph for 24 h, followed by a 6 h recovery in drug-free medium. RAD18 depletion was confirmed by Western blotting. Representative images are shown. Dot plot shows tail moment values. Horizontal lines indicate the median values (ns: $P > 0.05$; ***$P < 0.001$; ****$P < 0.0001$; Mann-Whitney test).

(**F**) **Effect of ATM inhibition on G4-FANCJ interactions detected by PLA.** Cells were pre-treated with 10 µM KU55933 (ATM inhibitor) for 1 h, followed by Aph treatment for 24 h ,with 10 µM MG132 during the last 4 h. PLA was performed using BG4 (G4) and anti-FANCJ antibodies. DNA was counterstained with DAPI. Dot plot shows the number of PLA spots per nucleus. Horizontal lines indicate median values (ns: $P > 0.05$; *$P < 0.05$; ****$P < 0.0001$; Kruskal-Wallis test).

**Figure S9. Inhibition of ATM phenocopies FANCJ deficiency**

**(A) Impact of ATM inhibition on FANCJ stability.** HeLa cells and FANCJ$^{KO}$ cells complemented with a phosphorylation-deficient mutant (FANCJ$^{KO+S990A}$) were pre-treated with 10 µM KU55933 (ATM inhibitor) for 1 h, treated or not with 24 h Aph and 10 µM MG132 during the last 4 h. Blots were probed with antibodies against FANCJ and LAMIN B1. Data are shown as super plot of

FANCJ/LAMIN B1 ratios normalized to control (ns: $P > 0.05$; **$P < 0.01$; ***$P < 0.001$; ****$P < 0.0001$; one-way ANOVA test).

**(B) Analysis of FANCJ stability in mutant cells**. FANCJ$^{KO+S990A}$ and FANCJ$^{KO+S990E}$ cells were treated or not with 40 µg/mL cycloheximide as indicated to assess FANCJ stability. Blots were probed with antibodies against FANCJ and GAPDH.

**(C) Effect of ATM inhibition on FANCJ phosphorylation status using a phospho-FANCJ Ser990 antibody.** HeLa and FANCJ$^{KO}$ cells were treated as in (A). Blots were probed with antibodies against anti-phospho-Ser990 FANCJ, total FANCJ and Topoisomerase IIβ.

**(D) DNA damage assessed by alkaline Comet assay.** Cells were treated as in (A). Representative images are shown. Dot plot shows tail moment values. Horizontal lines indicate median values (ns: $P > 0.05$; **$P < 0.01$; ****$P < 0.0001$; Mann-Whitney test).

**Figure S10. USP1 depletion promotes DNA damage accumulation**

**Assessment of DNA damage accumulation in USP1-depleted cells.** shWRNIP1$^{WT}$ and shWRNIP1$^{D37A}$ cells were depleted of USP1 by RNAi and, 48 h later, pre-treated with 10 µM KU55933 (ATM inhibitor) for 1 h, followed by Aph treatment for 24 h, with 10 µM MG132 during the last 4 h. Representative images are shown. Dot plot shows tail moment values. Horizontal lines indicate median values (ns: $P > 0.05$; *$P < 0.05$; **$P < 0.01$; Mann-Whitney test).
